# Activity-dependent mitochondrial transport in peri-synaptic glia drives motor function

**DOI:** 10.1101/2021.11.29.470476

**Authors:** Dunham D. Clark, Sonja A. Zolnoski, Emily L. Heckman, Michael R. Kann, Sarah D. Ackerman

**Affiliations:** Brain Immunology and Glia Center, Department of Pathology and Immunology, Washington University School of Medicine, Saint Louis, MO 63110; Institute of Neuroscience, Howard Hughes Medical Institute, University of Oregon, Eugene, OR 97403

**Author notes:** Current Address: Department of Microbiology, Perelman School of Medicine, University of Pennsylvania, Philadelphia, PA 19104. Current Address: Department of Molecular, Cellular, & Developmental Biology, University of Michigan, Ann Arbor, MI 48109. Current Address: University of Pittsburgh School of Medicine, Pittsburgh, PA 15213.

## Abstract

Neurons have an outsized metabolic demand, requiring continuous metabolic support from non-neuronal cells called glia. When this support fails, toxic metabolic byproducts accumulate, ultimately leading to excitotoxicity and neurodegeneration. Astrocytes, the primary synapse-associated glial cell type, are known to provide essential metabolites (*e.g.* lactate) to sustain neuronal function. Here, we leverage the well-characterized *Drosophila* motor circuit to investigate another means of astrocyte-to-neuron metabolic support: activity-dependent trafficking of astrocyte mitochondria. Following optogenetic activation, motor neuron mitochondria migrate away from synapses. By contrast, astrocytic mitochondria accumulated peri-synaptically, and at times, were transferred into neighboring neurons. A genetic screen identified the mitochondrial adaptor protein Milton as a key regulator of this process. Astrocyte-specific *milton* knockdown disrupted regular mitochondrial trafficking, resulting in locomotor deficits, dysfunctional motor activity, and altered synapse number at the neuromuscular junction. These findings suggest that astrocytes dynamically redistribute mitochondria to buffer metabolic demand at synapses, highlighting a potential mechanism by which glia protect neural circuits from metabolic failure and neurodegeneration.

## INTRODUCTION

Despite constituting just 2% of total body weight, the human brain accounts for ∼20% of the body’s day-to-day metabolic consumption (Annesley & Fisher, 2019; Wang et al., 2010). This extraordinary demand is a direct reflection of the metabolic requirements of neurons, which require a constant supply of energy to sustain signaling throughout their long lifespan (Balasubramanian, 2021). In the human brain, a single cortical neuron consumes ∼4.7 billion ATP molecules every second while at rest (Zhu et al., 2012). To generate ATP, cells mainly rely on two distinct but linked biochemical pathways: glycolysis and oxidative phosphorylation (OXPHOS) (Hall et al., 2012; Yates, 2024; Yellen, 2018). Glycolysis rapidly converts cytosolic glucose into pyruvate or, in anaerobic conditions, lactate, yielding a modest energy output of 2 ATP molecules per glucose (Chandel, 2021; Pellerin & Magistretti, 1994). OXPHOS, a mitochondrial pathway, further metabolizes the products of glycolysis to produce a significantly higher yield (>30 ATPs per glucose) at the cost of speed (Nath & Villadsen, 2015; Senior, 1988). Due to its higher efficiency, neurons primarily rely on OXPHOS to meet their bioenergetic needs (Hall et al., 2012; Yellen, 2018). However, OXPHOS alone is unable to provide enough ATP to fuel a frequently-firing neuron (Bélanger et al., 2011; Yellen, 2018). Instead, metabolic coupling with neighboring glial cells, particularly astrocytes, provides the rest (Bélanger et al., 2011a; Bonvento & Bolaños, 2021).

Astrocytes, the most abundant glial cell type in the central nervous system (CNS), are essential for neural circuit formation, synaptic regulation, and neurotransmitter recycling (Freeman, 2010; Kim & Chung, 2023). A single human astrocyte can interact with up to two million synapses simultaneously with their numerous, fine processes, allowing them to influence entire circuits down to individual synapses (Allen & Eroglu, 2017; Oberheim et al., 2009). Beyond regulation of synaptic signaling, astrocytes provide neurons with both structural and metabolic support (Allen & Eroglu, 2017; Bonvento & Bolaños, 2021). Unlike neurons, astrocytes primarily rely on glycolysis to meet their own bioenergetic demands, and to generate lactate that neurons can use to fuel OXPHOS in a pathway known as the astrocyte-neuron lactate shuttle (Bastian et al., 2022; Bélanger et al., 2011; Tsacopoulos et al., 1996). Despite their preference for glycolysis, high-resolution electron microscopy has revealed that a wide network of mitochondria are often localized within an astrocyte’s peri-synaptic processes (Kacerovsky et al., 2016), suggesting that these mitochondria may play an active role in synaptic maintenance and neuronal bioenergetic support (Aten et al., 2022). The cellular mechanisms that guide this peri-synaptic mitochondrial positioning remain undefined.

Mitochondria are highly dynamic organelles that undergo continuous transport, fission, and fusion to adapt to changes in the cellular environment (Berridge & Neuzil, 2017). Mitochondrial transport is mainly facilitated by microtubule-based motor proteins like Kinesin and Dynein, with assistance from actin-based transporters in certain contexts (Fung et al., 2023; Melkov et al., 2016; Melkov & Abdu, 2018). In neurons, mitochondrial trafficking is tightly regulated, ensuring localized ATP production and calcium buffering at sites of heightened metabolic demand (Birsa et al., 2013; Schwarz, 2013). Due to their unique morphology, neurons require a transport system for intracellular mitochondrial localization that is both robust and adaptable, allowing for rapid deployment and retrieval of mitochondria to and from distant processes in response to stimuli (Birsa et al., 2013; Schwarz, 2013). The extent to which astrocytes exhibit similar regulatory control over mitochondrial transport in response to external stimuli, neuronal activity for example, remains largely unexplored. Given their extensive synaptic interactions and role in metabolic support, the ability of astrocytes to reposition mitochondria dynamically, and the consequences of such repositioning, could define an additional layer of complexity to neuron-glia coupling.

Mitochondrial transfer from glia to neurons has been documented in various pathological contexts, such as after ischemic injury or neurodegenerative disease, where neurons struggling with metabolic insufficiency receive astrocyte-derived mitochondria to sustain function. (Annesley & Fisher, 2019; Berridge & Neuzil, 2017; Hayakawa et al., 2016; Zhou et al., 2024). Similarly, Müller glia, astrocyte-like cells in zebrafish, have been observed internalizing and degrading damaged neuronal mitochondria following oxidative stress (Hutto et al., 2023). While these cases suggest intercellular mitochondrial transport as a fail-safe mechanism, it remains unclear whether such transfer occurs under normal physiological conditions, and what role it may play in circuit development and function.

Using the *Drosophila* motor circuit, we sought to understand the relationship between neuronal activity and astrocyte mitochondrial dynamics. By leveraging complementary binary expression systems (*GAL4/UAS* and *LexA/LexAop*), we were able to simultaneously manipulate neuronal activity and directly visualize changes in the behavior of neuronal and astrocytic mitochondria. We found that astrocytes use both microtubule and actin-based molecular motors to re-position their mitochondria within the neuropil, and that this trafficking is necessary for long-term motor function. We hypothesize that this coordinated, activity-dependent relationship between astrocyte mitochondria and neurons allows these peri-synaptic glia to match metabolic need with local neuronal signaling and provide “energy on-demand” (Magistretti et al., 1999).

## RESULTS

### Activity-dependent positioning of astrocyte mitochondria

Astrocytes are an essential energy source for neurons, facilitating both neuronal longevity and long-term memory formation (Lago-Baldaia et al., 2020; Suzuki et al., 2011; Tsacopoulos & Magistretti, 1996). Consistent with their role in metabolic support, transmission electron microscopy has revealed mitochondria within the delicate peri-synaptic processes of astrocytes (Kacerovsky & Murai, 2016). However, whether these mitochondria accumulate through passive diffusion or active recruitment mechanisms remained unclear. To address this question, we utilized complementary binary expression systems (*Gal4/UAS, LexA/LexAop*) to label and manipulate the well-characterized aCC/RP2 motor neurons of the *Drosophila* larval motor circuit (Landgraf et al., 2003), and simultaneously label astrocytic mitochondria (Ackerman et al., 2021). First, we modified our previously-published image analysis pipelines in Imaris (Ackerman et al., 2021; Sales et al., 2019) to define pre-synapses (Brp+) that were co-localized with aCC/RP2 dendrites within individual hemisegments of the *Drosophila* ventral nerve cord (VNC, akin to the vertebrate spinal cord). In parallel, we reconstructed all astrocyte mitochondria within the same dendritic domain. Through a series of masking steps, we then determined (a) the percentage of these synapses that were associated with astrocyte mitochondria, alongside (b) the total volume of domain-associated mitochondria, (c) the total number of dendrite-associated mitochondria and (d) the average volume of each individual astrocytic mitochondria (Figure 1A, see Supplemental Figure 1 legend and methods for full details).

**Figure 1.**
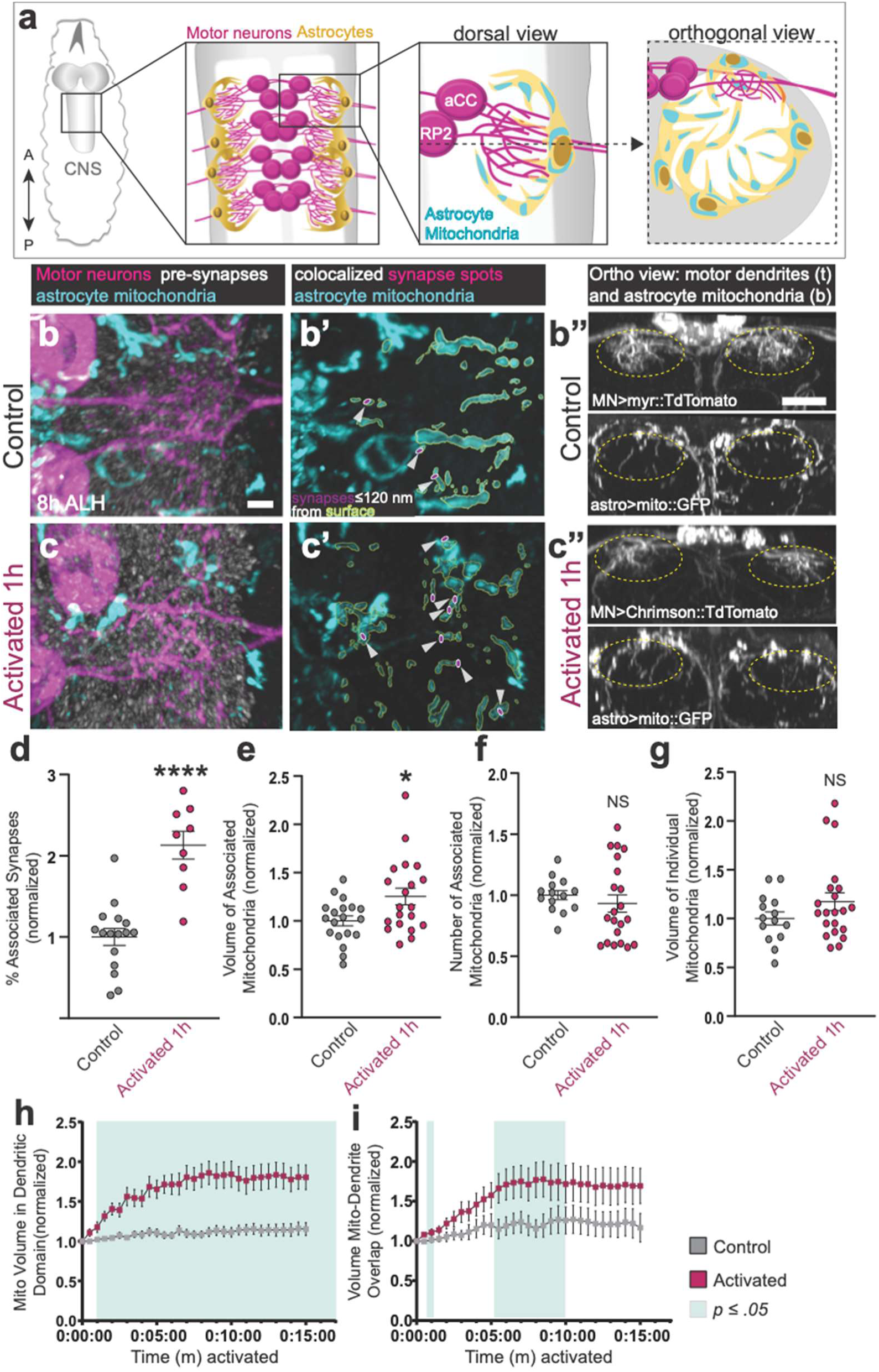
Motor neuron activation recruits astrocyte mitochondria to dendrites & synapses. **(a)** Schematic for reader orientation. A, anterior. P, posterior. Astrocyte (yellow) mitochondria (cyan) are distributed throughout the synapse-rich neuropil (white) near motor dendrites (magenta). **(b-c”)** Representative images of dark-reared control (**b-b”**) and 1-hour activated (**c-c”**) motor dendrites (magenta, *RN2-lexA, lexAop-Chrimson::TdTomato)* with neighboring astrocyte mitochondria (cyan, *R25H07-gal4, UAS-mito::GFP*) and presynapses (white, Brp+) in fixed preparations at 8 h ALH. Prime panels show synapse “Spots” co-localized (≤ 120 nm) with astrocyte mitochondria. Double prime panels show orthogonal views of motor dendrites (top, t) and astrocyte mitochondria (bottom, b) within the dendritic domain (yellow dashed circle). (**b-b’**, **c-c’**) Scale bar, 2 µm. (**b”**,**c”**) Scale bar, 10 µm. **(d)** Quantification of the percentage of synapses associated with astrocyte mitochondria (p<.0001, Mann-Whitney U test). Control: n=16 animals. Activated: n=9 animals. **(e)** Quantification of the mitochondrial volume in the dendritic domain (p<.05, Mann-Whitney test). Control: n=19 animals. Activated: n=20 animals. **(f)** Quantification of number of mitochondria in dendritic domain (p<.26, Mann-Whitney U test). Control: n=14 animals. Activated: n=21 animals. **(g)** Quantification of average volume of associated mitochondria in dendritic domain (p<.33, Mann-Whitney U test). Control: n=14 animals. Activated: n=21 animals. (**d-g**) Error bars = SEM. NS, not significant. N values reflect animals from 3 technical replicates. (**h-i**) Quantification of (**h**) astrocyte mitochondrial volume (*R25H07-gal4, UAS-mito::GFP*) within the dendritic domain and (**i**) overlapping mitochondria-dendrite volume in live samples (8 h ALH) imaged every 30 seconds for a 15-minute window of activation (magenta, *RN2-lexA, lexAop-Chrimson::TdTomato*) or control (grey, *RN2-lexA, lexAop-myr::TdTomato*). Control: n=14 animals. Activation: n=15 animals. Green areas denote timepoints when p<.05 by one-way ANOVA. Error bars = SEM. N values reflect animals from 2 technical replicates.

Using these model neurons, we previously characterized a critical period of motor circuit plasticity during development when motor neurons robustly change their dendrite structure in response to neuronal activity, peaking just before hatching and leveling off ∼8 hours after larval hatching (h ALH) (Ackerman et al., 2021). To assess activity-dependent mitochondria-synapse association without the confounds of structural plasticity, we therefore limited our studies to larval stages ≥ 8 h ALH, when the critical period is closed, and dendrite morphology is not dramatically affected by motor neuron activity. Following optogenetic activation of motor neurons for one hour (*RN2-lexA, lexAop-Chrimson::tdTomato*), we observed a redistribution of astrocyte mitochondria (*R25H07-gal4, UAS-mito::GFP*) from the periphery to the center of the dendritic domain (Figure 1b-c”). This resulted in a significant increase in total astrocyte mitochondrial volume within the dendritic domain and a significant increase in the percentage of motor synapses within close proximity (≤ 120 nm) to astrocyte mitochondria, though individual mitochondrial volume remained largely stable (Figure 1d-g).

Previous mammalian slice culture assays demonstrated that acute, pharmacological activation of neurons resulted in pausing of astrocyte mitochondria near synapses in a calcium-dependent manner (Jackson et al., 2014; Jackson & Robinson, 2015). To determine if *Drosophila* motor neuron activity causes astrocyte mitochondrial stalling at motor dendrites/synapses, we used live imaging to visualize real-time changes in mitochondrial behavior during optogenetic activation of neighboring motor neurons. In contrast to simple stalling of mitochondria at synapses, we observed rapid mobilization of astrocyte mitochondria from the periphery to the center of the dendritic domain within one minute of motor neuron activation, resulting in a significant increase in mitochondrial volume within the activated region. Interestingly, this volume plateaued within five minutes of stimulation, suggesting that *in vivo*, mitochondria are initially actively transported to dendrites/synapses, followed by stalling once a certain threshold of activity has passed (Figure 1h-i, Supplemental Movies 1-2).

We next tested whether optogenetic silencing of motor neurons (*RN2-gal4, UAS-GtACR2::EYFP*) alters astrocyte mitochondrial distribution (*alrm-lexA, lexAop-mcherry::mito.OMM*, Figure 2). We found that, while motor neuron activation was sufficient to recruit astrocyte mitochondria to synapses, the percentage of mitochondria-associated synapses was unchanged following motor neuron silencing (Figure 2a-c). We did observe a significant decrease in overall mitochondrial volume within the dendritic domain after silencing for one hour (Figure 2d), but not shorter lengths of time (Figure 2g-h, Supplemental Movies 3-4). Finally, silencing resulted in a decrease in the individual volume of astrocyte mitochondria, suggesting that the silencing of associated neurons may drive fission of astrocytic mitochondria (Figure 2f). In sum, astrocyte mitochondria traffic to areas of elevated neuronal activity and away from areas of low neuronal activity.

**Figure 2.**
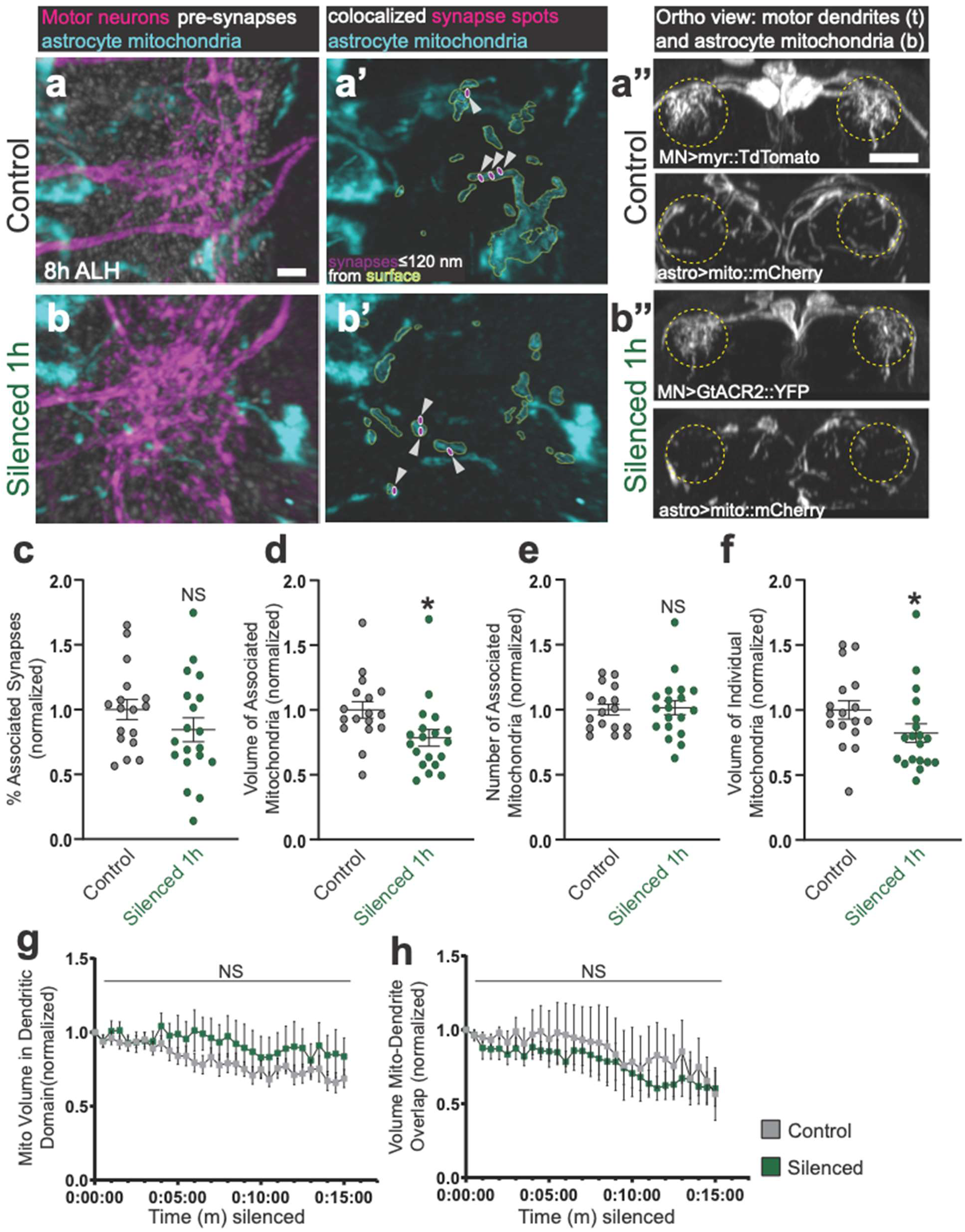
Motor neuron silencing repels astrocyte mitochondria from dendrites. **(a-b”)** Representative images of dark-reared control (**a-a”**) and 1-hour silenced (**b-b”**) motor dendrites (magenta, *RN2-gal4, UAS-GtACR2::EYFP)* with neighboring astrocyte mitochondria (cyan, *alrm-lexA, lexAop-mcherry::mito.OMM*) and presynapses (white, Brp+) in fixed preparations at 8 h ALH. Prime panels show synapse “Spots” co-localized (≤ 120 nm) with astrocyte mitochondria. Double prime panels show orthogonal views of motor dendrites (top, t) and astrocyte mitochondria (bottom, b) within the dendritic domain (yellow dashed circle). (**a-a’**, **b-b’**) Scale bar, 2 µm. (**a”**,**b”**) Scale bar, 10 µm. **(c)** Quantification of the percentage of synapses associated with astrocyte mitochondria (p<.26, Mann-Whitney U test). Control: n=17 animals. Silenced: n=19 animals. **(d)** Quantification of the mitochondrial volume in the dendritic domain (p<.003, Mann-Whitney U test). Control: n=17 animals. Silenced: n=19 animals. **(e)** Quantification of number of mitochondria in dendritic domain (p<.84, Mann-Whitney U test). Control: n=17 animals. Silenced: n=19 animals. **(f)** Quantification of average volume of associated mitochondria in dendritic domain (p<.03, Mann-Whitney U test). Control: n=17 animals. Silenced: n=19 animals. (**c-f**) Error bars = SEM. NS, not significant. N values reflect animals from 2 technical replicates. (**g-h**) Quantification of (**g**) astrocyte mitochondrial volume (*alrm-lexA, lexAop-mcherry::mito.OMM*) within the dendritic domain and (**h**) overlapping mitochondria-dendrite volume in live samples (8 h ALH) imaged every 30 seconds for a 15-minute window of silencing (green, *RN2-gal4, UAS-GtACR2::EYFP*) or control (grey, *RN2-gal4, UAS-myr::GFP*). Control: n=11 animals. Silencing: n=9 animals. Green areas denote timepoints when p<.05 by one-way ANOVA. Error bars = SEM. N values reflect animals from 2 technical replicates.

### Activity-dependent positioning of motor neuron mitochondria

The observation that astrocyte mitochondria rapidly reposition themselves toward active dendritic regions suggests a finely tuned mechanism by which glial cells sense and respond to the local energetic needs in neurons. As neurons possess their own mitochondrial networks essential for maintaining synaptic transmission and excitability (Sheng & Cai, 2012; Smith et al., 2016), it remained important to assess whether mitochondrial localization within motor neurons is likewise modulated by activity. We utilized the same optogenetic tools to activate or silence our model motor neurons (aCC/RP2). We then measured activity-dependent changes in motor neuron mitochondrial number, volume, and peri-synaptic location using the same analysis pipeline as in Figures 1-2, and Supplemental Figure 1 (see Methods for complete details).

Following optogenetic activation of motor neurons for one hour (*RN2-gal4, UAS-Chrimson::mVenus*), we observed a redistribution of neuronal mitochondria (*RN2-Gal4, UAS-mCherry.mito.OMM*) away from motor synapses (Figure 3a-b”). This resulted in a significant decrease in the percentage of motor synapses within close proximity (≤ 120 nm) to neuronal mitochondria (Figure 3c). While total mitochondrial volume within the dendritic domain did not change, we saw an activation-dependent decrease in the number of mitochondria and a corresponding increase in the volume of each individual mitochondria. These data suggest that mitochondrial fusion may be a driver of this re-localization (Figure 3e-f). Following optogenetic silencing of motor neurons for one hour (*RN2-gal4; UAS-GtACR2::YFP*), we observed a similar reduction in the number of neuronal mitochondria (*RN2-Gal4, UAS-mCherry.mito.OMM*) within the dendritic domain; however, we observed no changes in the percent of mitochondria-associated synapses or the volume of individual mitochondria (Figure 4). Collectively, these data suggest that motor neuron activity drives changes to the number of their own mitochondria. If this change in localization and number of mitochondria is linked, or even necessary, to the peri-synaptic localization of astrocytic mitochondria is unclear.

**Figure 3.**
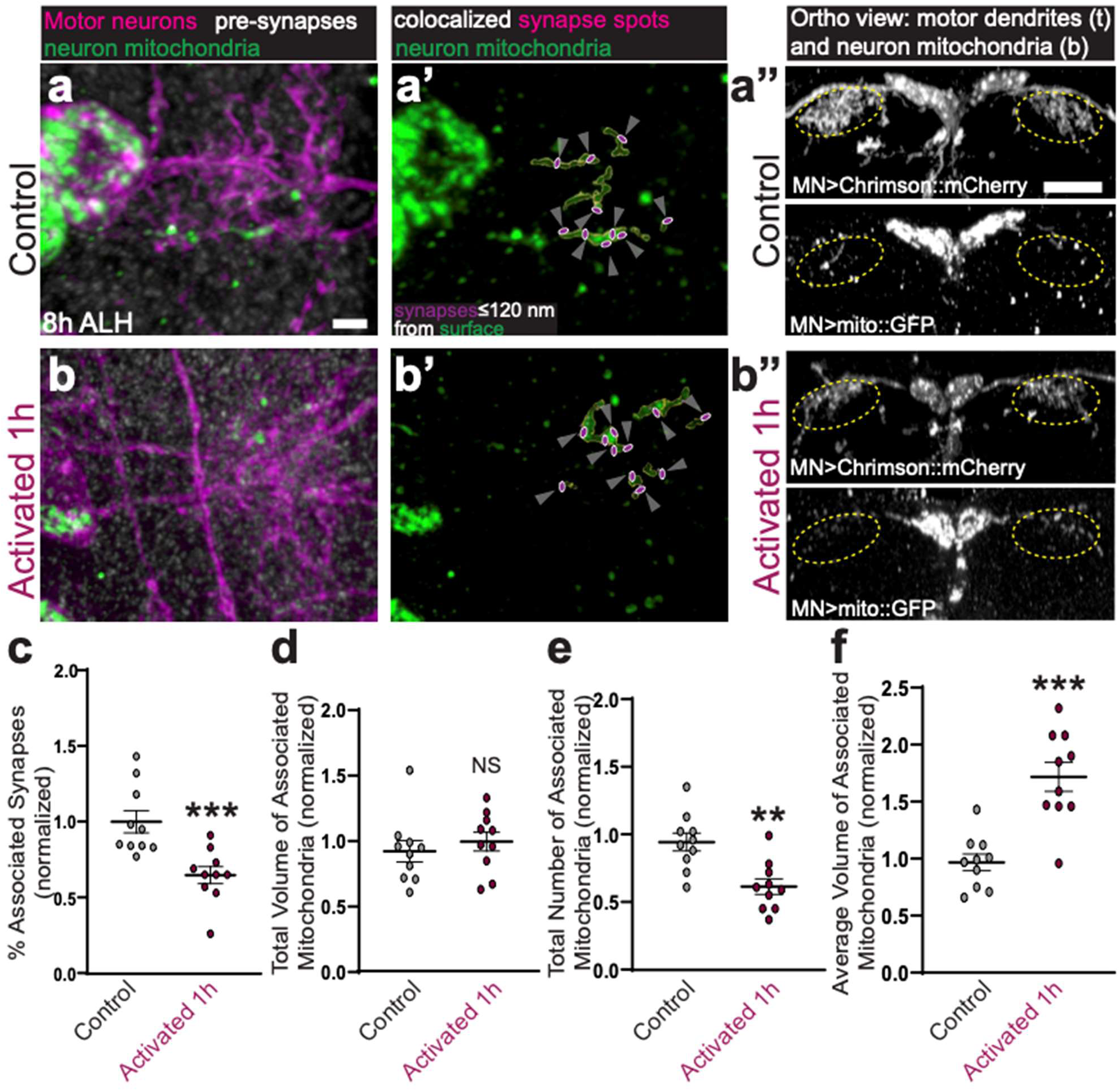
Motor neuron mitochondria flee synapses following activation. (**a-b”**) Representative images of dark-reared control (**a-a”**) and 1-hour activated (**b-b”**) motor dendrites (magenta, *RN2-gal4; UAS-Chrimson::mCherry*) with neuron mitochondria (green, *RN2-gal4, UAS-mito::GFP*) and presynapses (white, Brp+) in fixed preparations at 8 h ALH. Prime panels show synapse “Spots” co-localized (≤ 120 nm) with neuron mitochondria. Double prime panels show orthogonal views of motor dendrites (top, t) and neuron mitochondria (bottom, b) within the dendritic domain (yellow dashed circle). (**a-a’, b-b’**) Scale bar, 2 µm. (**a”,b”**) Scale bar, 10 µm. (c) Quantification of the percentage of synapses associated with neuron mitochondria (p=.0004, Mann-Whitney U test). Control: n=10 animals. Activated: n=10 animals. (d) Quantification of the mitochondrial volume in the dendritic domain (p=.2887, Mann-Whitney U test). Control: n=10 animals. Activated: n=10 animals. (e) Quantification of total number of associated mitochondria in the dendritic domain (p=.0020, Mann-Whitney U test). Control: n=10 animals. Activated: n=10 animals. (f) Quantification of the average volume (p=.0003, Mann-Whitney U test). Control: n=10 animals. Activated: n=10 animals. (**c-f**) Error bars = SEM. NS, not significant. N values reflect animals from 2 technical replicates.

**Figure 4.**
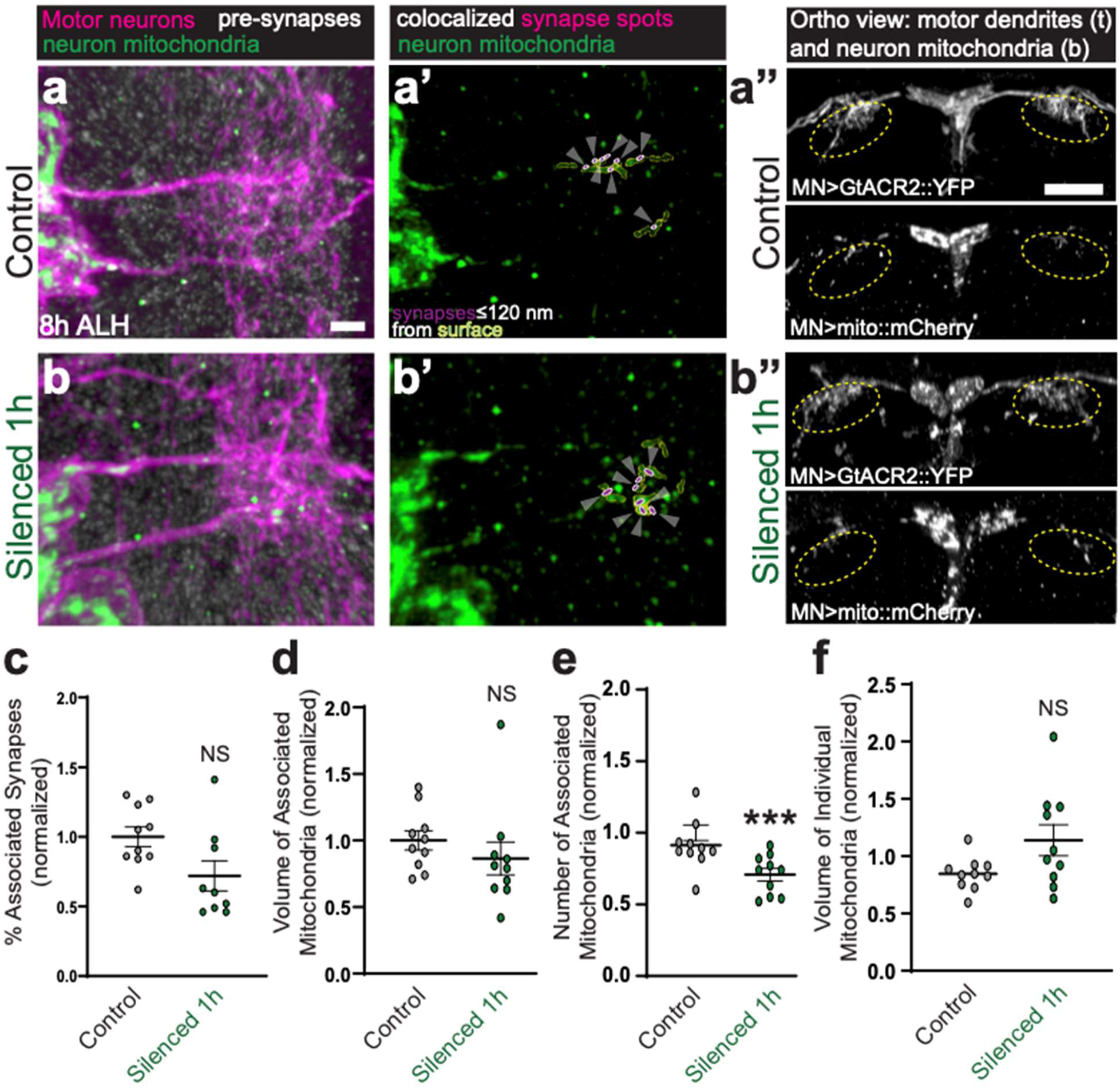
Motor neuron mitochondria flee synapses following silencing. **(a-b”)** Representative images of dark-reared control (**a-a”**) and 1-hour silenced (**b-b”**) motor dendrites (magenta, *RN2-gal4; UAS-GtACR2::YFP)* with neuron mitochondria (green, *RN2-gal4, UAS-mito::mCherry*) and presynapses (white, Brp+) in fixed preparations at 8 h ALH. Prime panels show synapse “Spots” co-localized (≤ 120 nm) with neuron mitochondria. Double prime panels show orthogonal views of motor dendrites (top) and neuron mitochondria (bottom) within the dendritic domain (yellow dashed circle). (**a-a’**, **b-b’**) Scale bar, 2 µm. (**a”**,**b”**) Scale bar, 10 µm. (c) Quantification of the percentage of synapses associated with neuron mitochondria (p=0.0506, Mann-Whitney U test). Control: n=10 animals. Silenced: n=9 animals. (d) Quantification of the mitochondrial volume in the dendritic domain (p=.0753, Mann-Whitney U test). Control: n=10 animals. Silenced: n=10 animals. (e) Quantification of the total number of associated mitochondria within the dendritic domain (p=.0004, Mann-Whitney U test). Control: n=10 animals. Silenced: n=10 animals. (f) Quantification of the average volume per associated mitochondria (p=.7959, Mann-Whitney U test). Control: n=10 animals. Silenced: n=10 animals. (**c-f**) Error bars = SEM. NS, not significant. N values reflect animals from 2 technical replicates.

### Astrocyte-to-neuron mitochondrial transport is suppressed by neuronal silencing

Thus far, we have shown that motor activity drives transport of astrocytic mitochondria towards motor synapses, whereas neuronal activity causes motor neuron mitochondria to retreat from the synapse. These data prompted us to ask: do motor neuron mitochondria evacuate the synapse to clear the way for astrocytic mitochondria to enter? To investigate whether neuronal activity induces astrocyte-to-neuron mitochondrial transfer, we used the *LexA/LexAop* and *Gal4/UAS* systems to specifically label and manipulate aCC/RP2 motor neurons (*RN2-lexA; lexAop-Chrimson::TdTomato*) alongside astrocytic mitochondria (*25H07-gal4;UAS-Mito::GFP*), as above (Figure 1), and we used an anti-GluL antibody to label astrocytic membranes (Theisen et al., preprint). We then used Imaris to assess whether astrocyte mitochondria could ever be seen inside motor neuron membranes. While we sometimes observed astrocyte mitochondria within motor dendrites at baseline (*e.g.* without motor neuron activation; Supplemental Figure 2a-a’’’), these events were rare. By contrast, we consistently observed astrocytic mitochondria within the cell bodies of adjacent motor neurons in activating and non-activating conditions. Accordingly, we developed an Imaris pipeline to mask motor neuron cell body membranes, astrocyte membranes, and astrocyte mitochondria to identify astrocyte-derived mitochondria enveloped within motor neuron cell bodies. We then quantified the following metrics within a standardized ROI: (1) colocalization of motor neuron signal with astrocyte mitochondria, (2) colocalization of motor neuron and astrocyte membranes, (3) colocalization of astrocyte membrane signal and astrocyte mitochondria, and (4) the number of astrocyte mitochondria fully enveloped within the motor neuron cell body (Figure 5a), see full details in legend and Methods). Under baseline conditions, astrocyte-derived mitochondria were largely confined to astrocytic processes, with some signal detected inside motor neurons. Following activation of motor neurons for 1 or 4 hours (see methods for details), astrocytic mitochondria were still observed within motor neuron cell bodies, but do not appear to be transferred at a rate higher than baseline conditions (Supplemental Figure 2). By contrast, motor neuron silencing for 4 hours suppressed intercellular trafficking of astrocyte-derived mitochondria into motor neuron cell bodies (Figure 5d-h). Together, these data suggest that astrocytes transfer mitochondria to neurons in response to basal activity, but that temporary increases in neuronal activity within a homeostatic range cannot potentiate this process.

**Figure 5.**
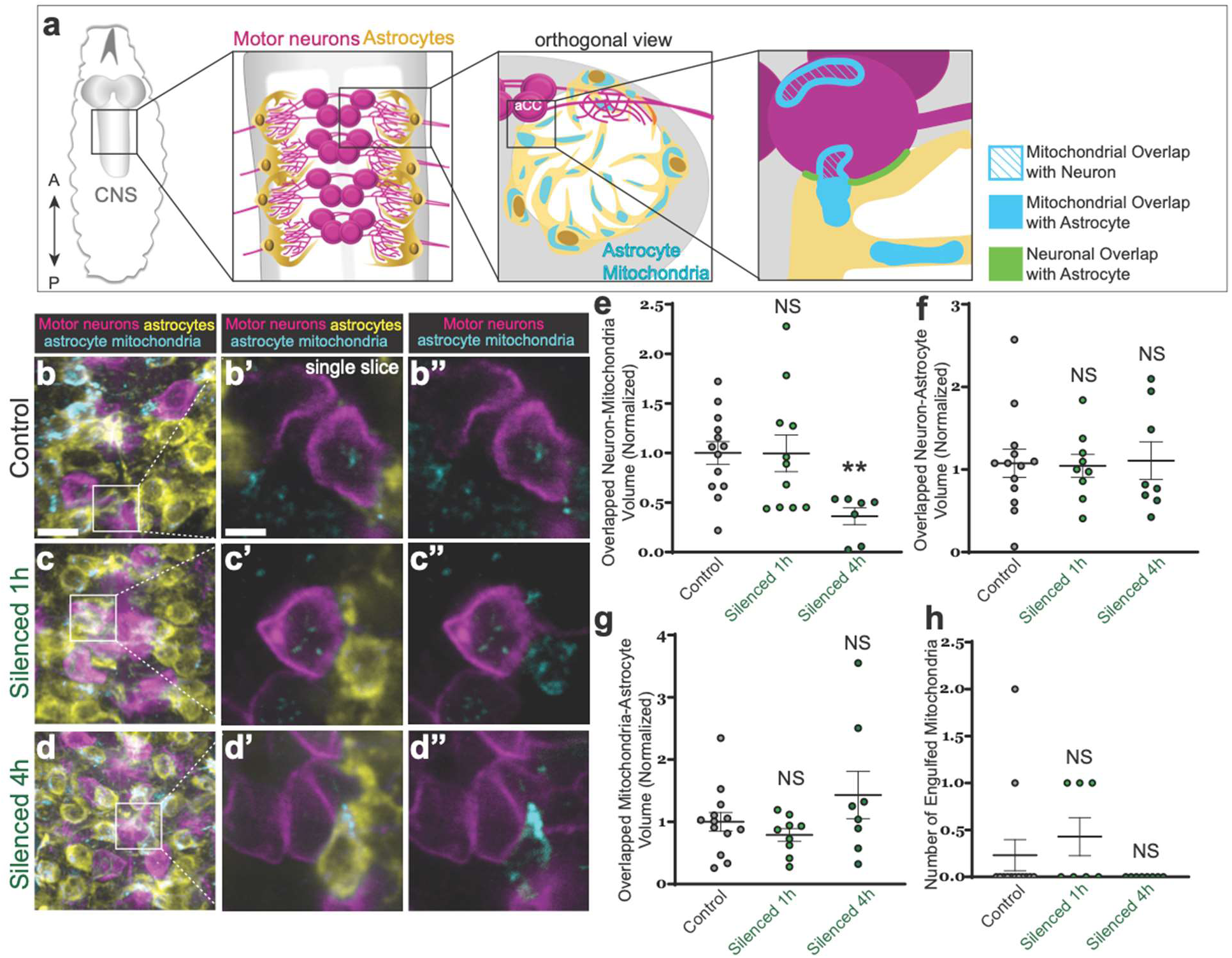
Intercellular transport from mitochondria from astrocytes to motor neuronss is impaired by neuronal silencing. (a) Schematic for reader orientation. A, anterior. P, posterior. Astrocyte (yellow) mitochondria (cyan) are distributed throughout the synapse-rich neuropil (white) near motor dendrites (magenta). (**b-d’**) Representative images of dark-reared control (**b-b’”**), 1-hour silenced (**c-c’’’**), and 4-hour silenced (**d-d’’’**) motor dendrites (magenta, *RN2-gal4, UAS-GtACR2::YFP*), astrocyte membranes (yellow, GLUL+) with astrocyte mitochondria (cyan, *alrm-lexA, lexAop-mito::mCherry*) in fixed preparations at 8 h ALH. Prime panels show a single 0.155 µm slice of astrocytes, neurons, and astrocyte mitochondria. Double prime panels show the same slice with motor neurons and astrocyte mitochondria. (**b,c,d**) Scale bar, 5 µm. Prime panels: Scale bar, 2 µm. (e) Quantification of the neuron-astrocyte mitochondrial overlap Control: n=16 animals. 1-hour silenced: n=11 animals ( p>0.9999, Kruskal-Wallis test). 4-hour silenced: n=7 (p=0.0087, Kruskal-Wallis test). (f) Quantification of astrocyte mitochondrial volume. Control: n=13 animals (p>0.9999, Kruskal-Wallis test). 1-hour silenced: n=9 animals (p>0.9999, Kruskal-Wallis test). 4-hour silenced: n=8 animals (p>0.9999, Kruskal-Wallis test). (g) Quantification of the neuron-astroctye membrane overlap. Control: n=13 animals. 1-hour silenced: n=9 animals (p>0.9999, Kruskal-Wallis test). 4-hour silenced: n=8 animals (p>0.9999, Kruskal-Wallis test). (**i**) Count of astrocyte mitochondria found completely enveloped by neuron membrane. Control: n=13 animals. 1-hour silenced: n=7 animals (p=0.5126, Kruskal-Wallis test). 4-hour silenced: n=8 animals (p>0.9999, Kruskal-Wallis test). (**e-h**) NS, not significant. Error bars = SEM. N values reflect animals from 2 technical replicates.

### Identification of cellular machinery for astrocyte mitochondrial trafficking

Having established a relationship between neuronal activity and the intracellular position of astrocytic mitochondria, we next aimed to define the mechanisms that facilitate this transport. In many diverse cell types, mitochondria are trafficked along cytoskeletal elements using molecular motor proteins that are specific to microtubules (*e.g.* Kinesins and Dyneins) or actin (*e.g.* Myosins) to reach distinct subcellular locations. Although kinesin-1 motors are the primary drivers of mitochondrial transport in neurons (Pilling et al., 2006), the process by which astrocytes traffic their mitochondria is vastly understudied. To define this mechanism in astrocytes, we performed astrocyte-specific RNA interference (RNAi) knockdown of molecular motor and mitochondrial adaptor proteins using previously published RNAi lines *(Perkins et al., 1997)*, and visualized astrocyte mitochondria (*R25H07-gal4, UAS-mito::GFP, UAS-RNAi*) by *in vivo* time-lapse imaging and in fixed preparations (Supplemental Movies 5-8, Figure 6). We screened ten lines targeting the adaptor protein *milton/TRAK1* (N=3 lines), the mitochondrial membrane GTPase *miro/RHOT1* (N=1 line, previously linked to astrocyte mitochondrial trafficking (Stephen et al., 2015)), the molecular motors *kinesin heavy chain* (N=2 lines), *dynein heavy chain* (N=2 lines), *didum/MYO5A* (N=1 line), and *jaguar/MYO6* (N=1 line). As above, we assessed defects in mitochondrial trafficking at 8 h ALH. During this developmental stage, astrocyte membranes are rapidly growing into the neuropil to tile in between neuronal synapses (Ackerman et al., 2021; Stork et al., 2014). We found that astrocyte-specific knockdown of *milton* completely blocked mitochondrial trafficking into the synapse-rich neuropil of both the central brain and ventral nerve cord, without any observable defects in the peri-synaptic localization of astrocyte processes (Supplemental Movie 6, Figure 6a-b”, quantified in h-i). We also observed significant reductions in mitochondrial trafficking following *miro*, *khc*, *didum,* or *jaguar* knockdown (Supplemental Movie 8, Figure 6c-g”, quantified in h-i).

**Figure 6.**
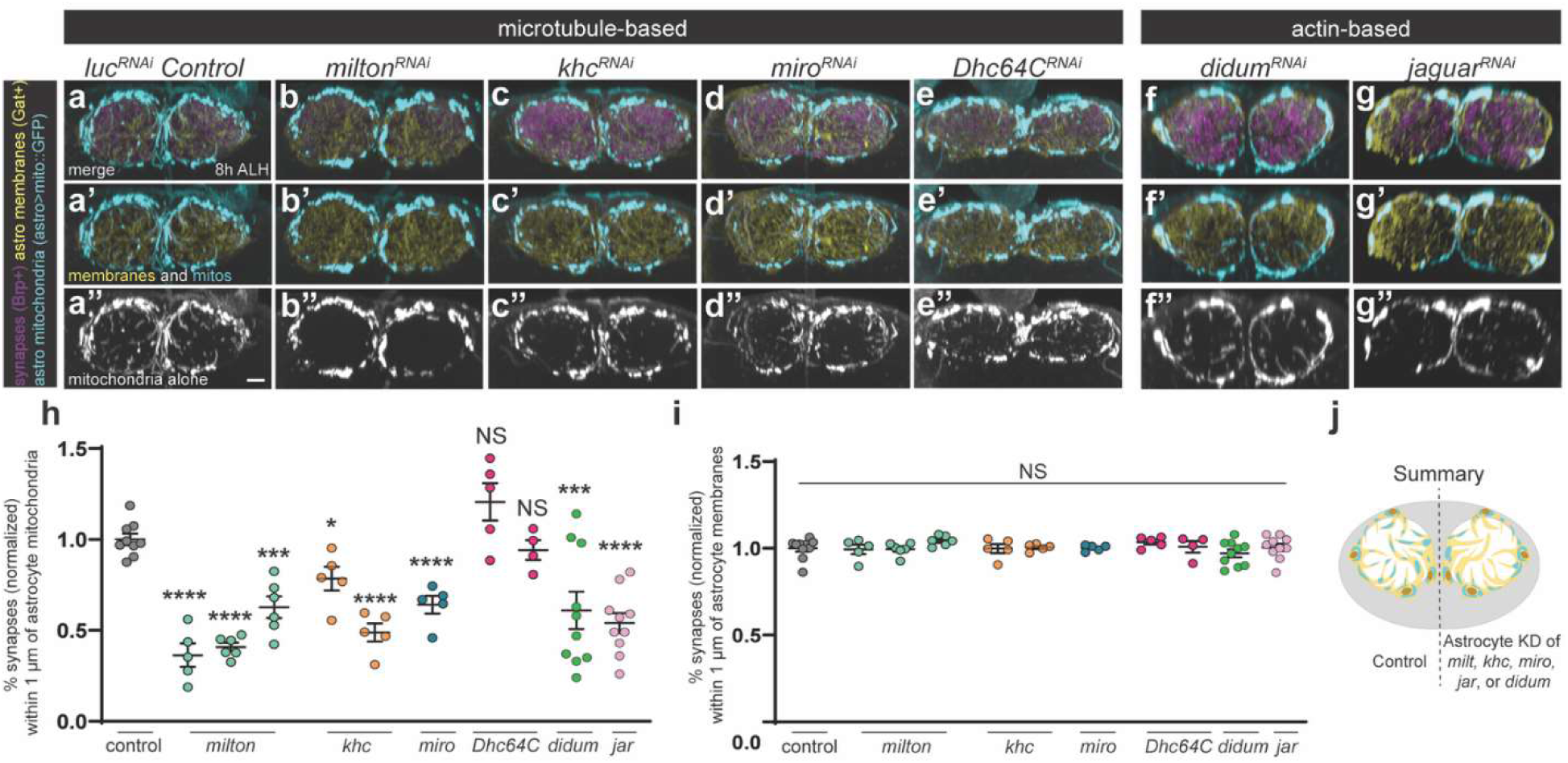
Identification of molecular machinery for astrocyte mitochondrial trafficking. **(a-g”)** Representative orthogonal views of astrocyte membranes (Gat+, yellow), mitochondria (*R25H07-gal4, UAS-mito::GFP*, cyan), and neighboring synapses (Brp+, magenta) in (**a-a”**) *luciferase^RNAi^* (luc) control brains at 8 h ALH, or (**b-g”**) 8 h ALH brains following astrocyte-specific knockdown (KD) of molecular machinery required for mitochondrial transport in axons. Prime panels show astrocyte membranes and mitochondria alone. Double prime panels show mitochondria alone. Scale bar, 5 µm for all panels. (h) Quantification of the percent of all synapses within 1 µm of astrocyte mitochondria. Statistical significance compared to control determined by one-way ANOVA: *milton^RNAi1^* (p<.0001), *milton^RNAi2^* (p<.0001), *milton^RNAi3^* (p<.002), *khc^RNAi1^* (p<.04), *khc^RNAi2^* (p<.007), *miro_RNAi_* (p<.0001), *Dhc64C^RNAi1^* (p<.14), *Dhc64C^RNAi2^* (p<.43), *didum^RNAi^* (p<0.0008), *jar^RNAi^* (p<0.0001). (i) Quantification of the percentage of all synapses within 1 µm of astrocyte membranes. Statistical significance compared to control determined by one-way ANOVA: *milton^RNAi1^* (p<.84), *milton^RNAi2^* (p<.85), *milton^RNAi3^* (p<.24), *khc^RNAi1^* (p<.91), *khc^RNAi2^* (p<.97), *miro^RNAi^* (p<.0001), *Dhc64C^RNAi1^* (p<.37), *Dhc64C^RNAi2^* (p<.88), *didum^RNAi^* (p0.9798), *jar^RNAi^* (p<0.9999). (**h-i**) Control: n=9. *milton^RNAi1^*: n=6. *milton^RNAi2^*: n=5. *milton^RNAi3^*: n=6. *khc^RNAi1^*: n=5. *khc^RNAi2^*: n=5 animals. *miro^RNAi^*: n=5 animals. *Dhc64C^RNAi1^*: n=5 animals. *Dhc64C^RNAi2^*: n=4 animals. *didum^RNAi^* (n=10), *jar^RNAi^* (n=10). Error bars = SEM. NS, not significant. N values reflect animals from 2 technical replicates. (j) Graphical summary of screen results.

In neurons, Milton is a mitochondrial adaptor protein that primarily serves as a linker between mitochondria-bound Miro and microtubule-associated Kinesin heavy chain (Glater et al., 2006; Schwarz, 2013). Together, this complex facilitates long-distance, (+) end-directed axonal trafficking of mitochondria along microtubules. While Milton also facilities mitochondrial trafficking along (-) end-directed microtubules (Glater et al., 2006), astrocyte knockdown of the (-) end-directed microtubule motor protein Dynein heavy chain did not have any impact on mitochondrial trafficking into the neuropil (Supplemental Movie 7, Figure 6e-e”, h). Finally, knockdown of actin-based motors (*jaguar*, *didum*) did not prevent mitochondrial trafficking entirely; however, both conditions, like knockdown of *milton, miro,* and *khc,* resulted in a significant reduction in mitochondria-synapse association without grossly affecting astrocyte morphogenesis (Figure 6f-j).

### Astrocyte mitochondrial trafficking deficits modify neuronal behavior and function

We defined Milton as a key mediator of astrocyte mitochondrial trafficking, prompting the question: what is the functional significance of astrocyte mitochondrial positioning in the neuropil? We proposed that peri-synaptic astrocyte mitochondria are essential for sustaining the high metabolic demands required for stable motor circuit structure and function. To address this hypothesis, we tested how loss of peri-synaptic astrocyte mitochondria influenced motor neuron activity (Ca^2+^) and motor-driven behaviors. Leveraging motor neuron-specific calcium imaging (*RN2-lexA, LexAop-GCaMP7s*), we found that perturbing astrocyte mitochondrial trafficking into the neuropil (*R25H07-gal4, UAS-milton^RNAi^*) caused a significant reduction in the frequency of Ca^2+^ spikes in aCC/RP2 motor neurons, although the amplitude of these spikes was largely unchanged (Figure 7a-c, Supplemental Movies 9-10). In parallel, we used a high-throughput behavior rig to assess motor output at both 8 h ALH and in late-stage larvae (larval instar 3, L3). Interestingly, while we observed only subtle behavioral differences at early larval stages (Supplemental Fig. 3), loss of astrocyte mitochondria from the neuropil resulted in significant locomotor deficits in late-stage larvae (L3), including reduced crawling speed and movement duration, lower total distance traveled, and increased bending frequency, indicative of compromised motor coordination (Figure 7g-k). Interestingly, astrocyte-specific knockdown of *milton* led to an increased number of synaptic boutons at the neuromuscular synapses downstream of aCC/RP2 at L3. These data suggest that impaired mitochondrial trafficking compromises motor neuron signaling in the CNS and may trigger compensatory, homeostatic plasticity in the periphery (Supplemental Figure 4).

**Figure 7.**
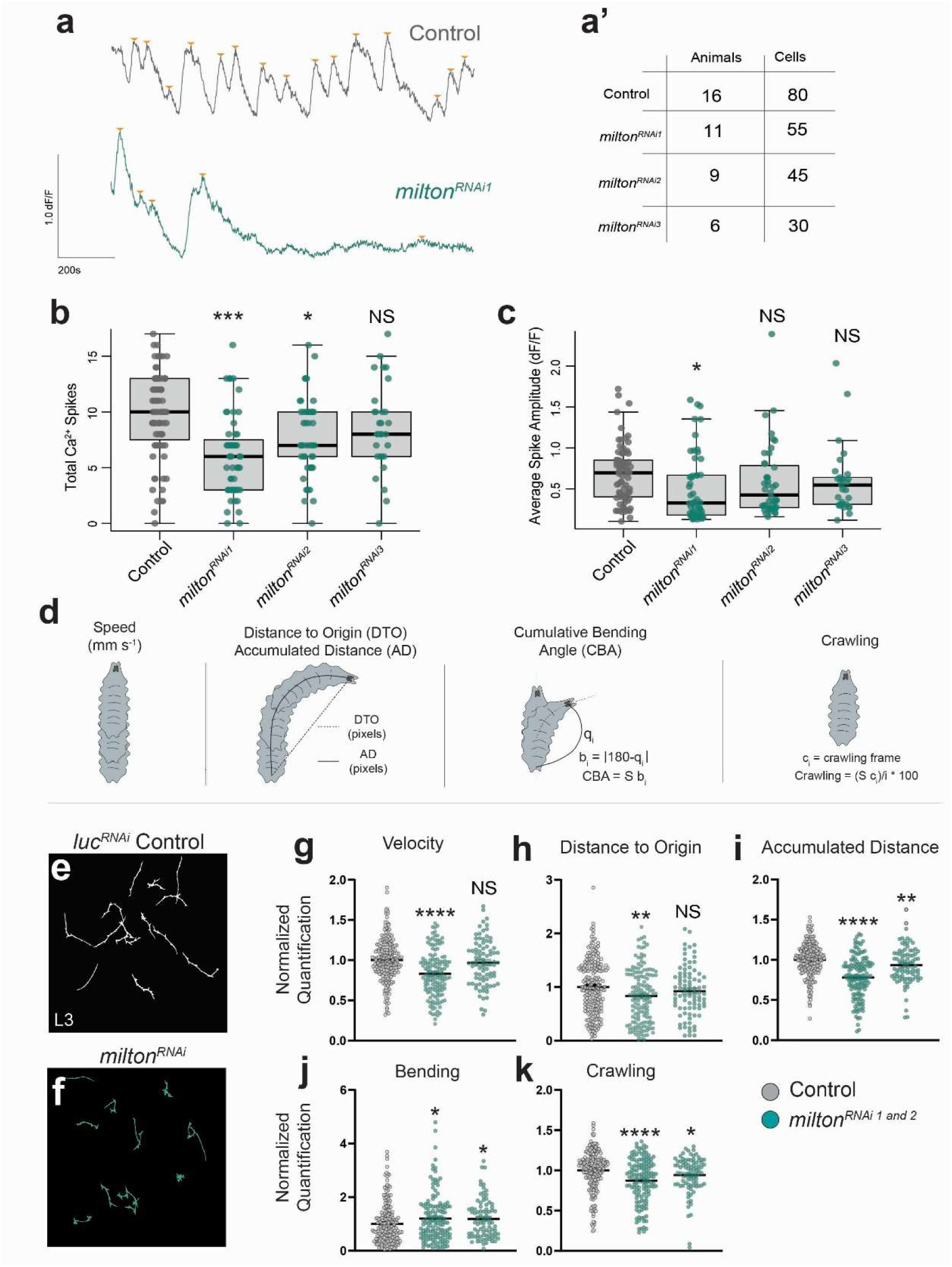
Astrocyte mitochondria are necessary for motor signaling and behavior. (a) Representative single cell dF/F traces from control (*luciferase^RNAi^*) and *milton^RNAi^* (*RN2-lexA, lexAop-GCaMP7s; 25H07-gal4,UAS-RNAi*) brain explants at 8 h ALH. Peaks of calcium (orange triangle) identified by PeakCaller. Prime panel shows the total number of animals and cells quantified for **b-c**. (b) Frequency of calcium peak events for control or following astrocyte *milton* knockdown with three independent RNAi lines. Each point represents an individual cell. Control: n=80 cells from 16 animals, *milton^RNAi1^*: n=55 cells from 11 animals (p<.0001, Student’s t-test), *milton^RNAi2^*: n=45 cells from n=9 animals (p<.006, Student’s t-test). *milton^RNAi3^*: n=30 cells from 6 animals (p=0.152, Student’s t-test). Error bars, SD. NS, not significant. (c) Average spike amplitude for control or following astrocyte *milton* knockdown with three independent RNAi lines. Each point represents an individual cell. Control: n=80 cells from 16 animals, *milton^RNAi1^*: n=55 cells from 11 animals (p<.01, Student’s t-test), *milton^RNAi2^*: n=45 cells from n=9 animals (p<.205, Student’s t-test). *milton^RNAi3^*: n=30 cells from 6 animals (p=0.296, Student’s t-test). Error bars, SD. NS, not significant. (d) Schematic of behavioral paradigms tested at larval stage 3 (L3). **(e-f)** FIMTrack traces from individual, free-crawling larvae recorded for 1 minute at 4 Hz*. luciferase^RNAi^* (luc) Control: n=222 animals. *milton^RNAi1^*: n=139 animals. *milton^RNAi2^*: n=85 animals. N values reflect animals from 3 technical replicates. **(g-k)** Quantification of behavioral metrics relative to control. Statistical significance determined by Mann-Whitney U Test for (**f**) velocity (*milton^RNAi1^*, p<.0001; *milton^RNAi2^*, p<.37), (**g**) distance to origin (*milton^RNAi1^*, p<.01; *milton*^RNAi2^, p<.25), (**h**) accumulated distance (*milton^RNAi1^*, p<.0001; *milton^RNAi2^*, p<.01), **(i)** cumulative bending angle (*milton^RNAi1^*, p<.04; *milton^RNAi2^*, p<.02), and (**j**) crawling (*milton^RNAi1^*, p<.0001; *milton^RNAi2^*, p<.05). Error bars = SEM. NS, not significant.

## DISCUSSION

### Metabolic demand of neurons requires external protection against neurodegeneration

Sustained neuronal signaling relies primarily on mitochondrial oxidative phosphorylation (OXPHOS). OXPHOS produces far more ATP than glycolysis, but inevitably generates reactive oxygen species (ROS) as byproducts (Lardy & Ferguson, 1969; Reczek & Chandel, 2015), which ultimately drive DNA mutagenesis and mitochondrial dysfunction (Reczek & Chandel, 2015). Neurons are uniquely vulnerable to these insults: as terminally differentiated, post-mitotic cells, they are unable to dilute or replace damaged mitochondria through division (Harman, 1972). Thus, the prevailing hypothesis is that neurons rely on metabolites from neighboring glial cells to sustain firing while limiting oxidative consequences.

### Neuronal activity dynamically regulates astrocyte mitochondrial position

Here, we identify one mechanism by which astrocytes may metabolically support neurons— through dynamic relocalization of their mitochondria. TEM data previously indicated that peri-synaptic astrocytic processes are enriched for mitochondria (Kacerovsky & Murai, 2016). Whether this localization is required, potentially for astrocytes to metabolically support their neighbors, or to otherwise promote neuronal health in the local environment, remained unclear (Bélanger et al., 2011). Using live imaging, we observed a near-immediate, activity-dependent increase in astrocyte mitochondrial mobility, until the local population stalled near firing neurons. We observed the reverse upon prolonged silencing, indicating that these organelles are dynamically recruited and withdrawn in accordance with local levels of activity. This behavior parallels findings in murine hippocampal slice cultures, where neuronal stimulation induced calcium-dependent mitochondrial stalling within astrocytic processes (Jackson et al., 2014), pointing to a conserved signaling axis linking synaptic activity to subcellular mitochondrial placement.

### Milton-based mitochondrial transport in astrocytes is required for motor function

In neurons, the machinery required to facilitate subcellular mitochondrial location is well-characterized, yet the mechanisms governing the same phenomena in astrocytes is understudied (Glater et al., 2006). Neuronal mitochondrial transport relies heavily on adaptor proteins such as Milton (TRAK1), which link mitochondria to molecular motors including Kinesin and Dynein, facilitating their movement along microtubules (Schwarz, 2013). Our data also identify Milton as essential for mitochondrial trafficking within astrocytes; astrocyte-specific knockdown of *milton* profoundly impaired mitochondrial transport toward synapse-rich regions without altering the overall morphology of the glial cells. Additionally, astrocyte-specific knockdown of myosin motors *jaguar* (MYO6) and *didum* (MYO5A) similarly disrupted mitochondrial trafficking within the synapse-rich neuropil. These data demonstrate that, as with neurons (Course et al., 2016), actin-based transport mechanisms work alongside the canonical microtubule-based system to facilitate trafficking of mitochondria in astrocytes (Glater et al., 2006). Disrupting mitochondrial trafficking in astrocytes through astrocyte-specific knockdown of *milton* was sufficient to impair synapse-associated mitochondrial coverage, alter endogenous neuronal activity, and disrupt coordinated locomotor behavior. This phenotype, emerging in the absence of exogenous stressors or injury, underscores the essential role of astrocytic peri-synaptic mitochondria in sustaining neuronal function. These findings mirror those in neurons across multiple model systems, where Milton/TRAK1 is essential for mitochondrial mobility and synaptic function, and whose dysfunction is linked to neurodevelopmental and neurodegenerative phenotypes (Brickley & Stephenson, 2011; Schwarz, 2013).

### Intercellular transport of mitochondria from astrocytes can be modulated by neuronal activity

In the context of injury, disease, or extreme metabolic stress, various non-neuronal cells have been shown to donate mitochondria to compromised neurons to sustain neuronal function and viability (Hayakawa et al., 2016; Joshi et al., 2019; Tseng et al., 2021). The transfer of mitochondria from astrocytes to neurons *in vivo*, during homeostatic conditions, had not been assayed. We observed basal transfer of astrocyte mitochondria into neighboring motor neuron cell bodies, and occasionally, into motor dendrites (Supplemental Figure 2a). Interestingly, this could not be potentiated by boosting neuronal activity within a physiological range (Supplemental Figure 2). These data suggest that astrocyte-to-neuronal transport may happen at low levels to sustain normal neuronal activity, but that high levels of intercellular transport are only engaged under pathological conditions (Hayakawa et al., 2016; Joshi et al., 2019; Tseng et al., 2021). Accordingly, neuronal silencing was able to suppress transfer beneath this basal level (Figure 5e).

In addition to intercellular transport of mitochondria from glia to neurons, a recent report showed that in zebrafish, mitochondrial stress (ROS) in cone photoreceptors can instead drive transfer of pathological mitochondria from neurons into neighboring astrocyte-like Müller glia for transmitophagy. Whether mitochondrial transfer from neurons to glia occurs in *Drosophila*, in pathological or physiological conditions, remains to be tested. Overall, our study identifies astrocyte peri-synaptic mitochondria as essential for sustained motor neuron function and motor-driven behaviors.

## Author Contributions

SDA conceived of the project; SDA performed and analyzed experiments in Figures 1-2, 6-7, and Supplemental Figures 3-4. DDC performed and analyzed experiments in Figures 3-6, and Supplemental Figure 2. SDA and SAZ performed and analyzed experiments in Figure 7 and Supplemental Figure 4. ELH analyzed GCaMP7s data in Figure 7. MRK analyzed behavior data in Supplemental Figure 3. DDC, SDA, SAZ, and ELH wrote the paper and prepared the figures. All authors commented and approved of the manuscript. We also thank Dr. Claudia Han for insightful feedback on the manuscript. This research was initially supported by a Milton Safenowitz postdoctoral fellowship to SDA, and was completed with the combined support of a BRAIN Initiative K99/R00 (NS121137) and support of Washington University School of Medicine startup funds to SDA.

## Competing Interest Statement

The authors declare no competing financial or non-financial interests.

## Data Availability Statement

This study did not generate code. Raw data for any main or supplemental figure will be deposited into Mendeley Data upon publication.

## Methods

### Lead Contact and Materials Availability

Additional information and inquiries regarding resources and reagents should be directed to and will be fulfilled upon request by Lead Contact Sarah D Ackerman.

### Experimental Model and Subject Details

#### Fly husbandry

We raised all fly stocks at 25°C on standard cornmeal fly food. Animals were staged relative to a 25°C standard. At 25°C, embryos take 21 hours to hatch into larvae (Crisp et al., 2008); larvae hatch and develop 1.25x faster at 30°C, and 2x slower at 18°C.

Transgenes (in order of appearance)

1. *RN2-lexA* (Ackerman et al., 2021)
2. *13lexAop-CsChrimson::tdTomato* (Ackerman et al., 2021)
3. *R25H07-gal4* (BDSC# 49145)
4. *UAS-mito-HA-GFP/cyo* (BDSC# 8442)
5. *13xLexAop2-CD4-tdTom/T6b* (BDSC# 77139)
6. *RN2-gal4* (Landgraf et al., 2003)
7. *UAS-GtACR2::EYFP* (Mauss et al., 2017)
8. *alrm-lexA::GAD* (Chromosome 3) (*Doherty et al., 2009*)
9. *lexAop-mCherry::mito.OMM* (BDSC# 66530)
10. *10XUAS-myr::GFP* (BDSC# 32198)
11. *UAS-mito-HA-GFP* (chromosome 3, BDSC# 8443)
12. *UAS-luciferase*^RNAi^ (BDSC# 31603)
13. *alrm-gal4* (Chromosome 2) (*Doherty et al., 2009*)
14. *10XUAS-mcd8::GFP* (BDSC# 5137)
15. *UAS-milton*^RNAi^ (BDSC#s 28385, 43173, 44477)
16. *UAS-khc*^RNAi^ (BDSC#s 35409, 35770)
17. *UAS-miro*^RNAi^ (BDSC# 43973)
18. *UAS-Dhc64C*^RNAi^ (BDSC# 36583, 36698)
19. *UAS-jaguar*^RNAi^ (BDSC# 28064)
20. *UAS-didum*^RNAi^ (BDSC# 55740)
21. *13XLexAop-IVS-jGCaMP7s* (BDSC# 80913)
22. *Mi{Trojan-lexA:QFAD.2}VGlut[MI04979-TlexA:QFAD.2]/CyO* (BDSC# 60314)
23. *13XLexAop2-IVS-myr::GFP* (X Chromosome, BDSC# 32211)
24. *13XLexAop2-IVS-CsChrimson::mVenus* (BDSC# 55139)

#### Animal collections

*For imaging of brains without optogenetics* (Figures 1-5, Supplemental Movies 1, 3, 5-10): We reared crosses at 25°C in collection bottles fitted with 3.0% agar apple juice caps seeded with plain yeast paste (Baker’s yeast + water). We then collected embryos on fresh 3.0% agar apple juice caps seeded with plain yeast paste for 1.5 hours (h) and aged at 25°C until hatching. At 0 h after larval hatching (ALH), larvae were collected and transferred to fresh apple caps (N=20 larvae per cap) containing yeast paste and aged until 8 h ALH, when they were dissected. For live imaging experiments, larvae were dissected at 8 h ALH and transferred to the confocal microscope for manipulation/recording (read more below).

*For imaging of brains with optogenetics* (Figures 1-5, Supplemental Movies 2, 4): We reared crosses at 25°C in collection bottles fitted with 3.0% agar apple juice caps seeded with fresh yeast paste that was supplemented with 0.5mM all-*trans* retinal (+ATR) (Sigma-Aldrich, R2500-100MG). Crosses were supplied fresh yeast paste daily (+ATR) for a minimum of 72 h prior to embryo collection to ensure maternal transfer of ATR into embryos. We then collected embryos on 3.0% agar apple juice caps with yeast paste (+ATR) for 1.5 h and aged at 25°C. To prevent premature optogenetic activation, we kept all crosses, embryos, and larvae in dark conditions until the appropriate developmental stage. At 25° C, *Drosophila* embryos hatch at 21 h after egg laying (AEL). Larvae were collected at 0 h ALH and transferred to fresh apple caps (N=10 larvae per cap) containing yeast paste (+ATR). For fixed preparations, larvae were then transferred to activating/silencing conditions for 1 hour and dissected at 8 h ALH. For live imaging experiments, larvae were dissected at 8 h ALH and transferred to the confocal microscope for manipulation/recording (read more below).

*For imaging of RNAi brains* (Figures 6-7, Supplemental Figure 4): We reared crosses at 25°C in collection bottles fitted with 3.0% agar apple juice caps seeded with plain yeast paste (Baker’s yeast + water). We then collected embryos on fresh 3.0% agar apple juice caps seeded with plain yeast paste for 1.5 h and aged at 30°C until hatching to maximize RNAi expression. For assessment of mitochondrial trafficking, larvae were collected at 0 h ALH and transferred to apple caps (N=20 larvae per cap) containing fresh yeast paste, aged 6.5 h (∼ 8 h ALH at 25°C standard), when they were dissected and prepared for imaging. For assessment of synapse number at the neuromuscular junction, apple caps were transferred to fresh food bottles (corn meal) when animals reached 0 h ALH. Bottles were maintained at 30°C for 60 h (∼ 72 h ALH at 25°C standard), when larvae were collected, dissected, and prepared for imaging.

*For behavioral analyses* (Figure 7, Supplemental Figure 3): We reared crosses at 25°C in collection bottles fitted with 3.0% agar apple juice caps containing plain yeast paste. Embryos were then collected on 3.0% agar apple juice caps with plain yeast paste for 1.5 h and aged at 30°C. For 8 h ALH behavioral analyses, apple caps were maintained at 30°C on the collection cap for 6.5 h (∼ 8 h ALH at 25°C standard) and then transferred to the behavioral rig for testing. For 3^rd^ instar stage behavioral analyses, after collection, apple caps were transferred to fresh food bottles (corn meal). Bottles were maintained at 30°C for 77 h (∼ 96 h ALH at 25°C standard) until larvae reached wandering 3^rd^ instar stage, when they were transferred to the behavioral rig for testing.

## METHOD DETAILS

### Sustained optogenetic activation/silencing

We used the following published strategy (Ackerman et al., 2021) for optogenetic activation/silencing in the following fixed preparation experiments (Figures 1-2). At the designated stage, dark-reared larvae (maintained on apple caps with yeast + ATR) were placed beneath a full spectrum light bulb, shining with an average intensity of 10550 lx (determined with two independent photometer software programs developed for Android: Light Meter© and Photometer©). We then dissected larvae in low-light conditions (<100 lx) immediately following activity manipulations. Yeast paste was placed in the middle of the apple cap to motivate larvae to remain on the top of the agar plate throughout the duration of the manipulation. Any larvae found outside of this area were not processed further.

### Activation/Silencing with a frequency-controlled “OptoChamber”

We used the following strategy for optogenetic activation/silencing in the following fixed preparation experiments (Figure 3-5, Supplemental Figure 2). At the designated stage, dark-reared larvae (maintained on apple caps with yeast) were placed within our custom designed, programable “OptoChamber” (Kann et al., 2024). This “OptoChamber” was then programmed to stimulate optogenetic constructs at peak wavelength and frequency, varying for each construct. Chrimson is optimally activated at 590 nm with minimal sensitivity below 500 nm (Klapoetke et al., 2014), while GtACR2 peaks at 470 nm and is largely inactive at ≥500 nm (Govorunova et al., 2015). The “OptoChamber” was programmed to deliver 590 nm or 470 nm light at 10 Hz for one hour, or no light (dark-reared control), using RGB LEDs rated at 600 mcd (red), 1000 mcd (green), and 500 mcd (blue). Larvae received either 1 (Figures 3-5, S2) or 4 (Figure 5, S2) hours of light treatment and were then immediately dissected following the same protocol as listed under “Sustained optogenetic activation/silencing”.

### Immunohistochemistry

Larval brains were dissected in sterile-filtered, ice-cold 1X PBS and mounted on 12mm #1 thickness poly-D-lysine coated round coverslips (Corning® BioCoat™, 354085 or German Glass Cover Slips, Fisher Scientific, 72196-12). We fixed brains for 12 minutes in fresh 4% paraformaldehyde (Electron Microscopy Sciences, 15710) in .3% PBSTriton, and then washed in .3% PBSTriton to remove fixative. For larval fillet preps, we dissected larvae in sterile-filtered, ice-cold HL3.1, fixed for 45 minutes in fresh 4% paraformaldehyde in detergent-free 1X PBS (Electron Microscopy Sciences, 15710), and then washed in .3% PBSTriton to remove fixative. At this point, all samples were treated the same way through post-secondary washes. Samples were blocked overnight at 4°C in .3% PBSTriton supplemented with 1% BSA (Fisher, BP1600-100), 1% normal donkey serum and 1% normal goat serum (Jackson ImmunoResearch Laboratories, Inc., 017-000-121 and 005-000-121). Samples were then incubated in primary antibody for one-two days at 4°C. The primary was removed, and brains were washed overnight at 4°C with 0.3% PBST. Samples were then incubated in secondary antibodies overnight at 4°C. The secondary antibodies were removed, and samples transferred to .3% PBSTriton overnight. <u>Larval brains</u> were then prepared for DPX mounting. Brains were dehydrated with an ethanol series: 30%, 50%, 70%, 90%, each for 5 minutes, then twice in 100% ethanol for 10 minutes each (Decon Labs, Inc., 2716GEA). Finally, samples were incubated in xylenes (Fisher Chemical, X5-1) for 2 x 10 minutes, were mounted onto slides containing DPX mountant (Millipore Sigma, 06552), and cured for 1-2 days before imaging. After post-secondary .3% PBST washes, <u>larval fillet preps</u> were washed into 1X PBS overnight. Fillet preps were then incubated in Phalloidin-405 (Alexa Fluor™ Plus 405 Phalloidin, a30104) in 1X PBS for 5 minutes, washed back into 1X PBS, and mounted in ProLong Glass Antifade Mountant (Thermo Fisher Scientific, P36982).

The following primary and secondary antibodies were used:

| Primary Antibody<br>(concentration) | Source | Figures |
| --- | --- | --- |
| Chicken anti-GFP (1:1000) | Aves Cat. GFP-1010 | 1-6, S1,<br>S2, S4 |
| Rabbit anti-cherry (1:500) | Novus Biologicals, Cat. NBP2-25157 | 2-4 |
| Mouse anti-Brp/Nc82 (1:100) | DSHB nc82 | 1-4, 6,<br>S1 |
| Rabbit anti-dsred (1:500) | Takara Bio Cat. 632496 | 1, S1,<br>S2 |
| Rat anti-cherry (1:500) | Fisher, M11217 | 5 |
| Mouse Anti-GLUL (1:400) | DSHB (BCN148.1.2H5) | 5, S2 |
| Rabbit anti-Gat (1:400) | M. Freeman lab | 6 |
| Ms anti-Dlg (1:100) | DSHB 4F3 | S4 |
| Phalloidin-405 (1:40) | Alexa Fluor™, a30104 | S4 |

Secondary Antibodies: All secondary antibodies were purchased from Jackson ImmunoResearch and used at a working concentration of 1:400. The following antibodies were used: Alexa Fluor® Rhodamine RedTM-X Donkey-Anti Mouse (715-295-151), Alexa Fluor® 647 AffiniPure Donkey Anti-Mouse IgG (715-605-151), Alexa Fluor® 488 Donkey anti-Chicken (703-545-155), Alexa Fluor® 647 Donkey-Anti Rabbit (711-605-152); Alexa Fluor® 488 Donkey anti-Rabbit (711-545-152), Alexa Fluor® Rhodamine RedTM-X Donkey Anti-Rabbit (711-295-152).

### Light Microscopy

*For fixed preparations:* Fixed larval preparations for synapse quantifications (central and NMJ: Figures 1-6, S4) were imaged on either a Zeiss LSM 800 laser scanning confocal fitted with a 63x/1.40 NA Oil Plan-Apochromat DIC m27 objective lens and GaAsP photomultiplier tubes or on a spinning disc confocal from Intelligent Imaging Innovations (3i) using a 63x/1.4 NA Oil Plan-Apochromat DIC m27 objective lens. Fixed larval preparations for astrocyte morphology analyses (Figure 6) were imaged on either a Zeiss LSM 700 laser scanning confocal using a 63x/1.4 NA Oil Plan-Apochromat DIC m27 objective lens (for microtubule-based motor proteins) or on a spinning disc confocal from Intelligent Imaging Innovations (3i) using a 63x/1.4 NA Oil Plan-Apochromat DIC m27 objective lens (for actin-based motor proteins). Fixed larval preparations for motor neuron mitochondria (Figures 3-5, S2) and intercellular mitochondrial transport were imaged on a custom-built spinning disc confocal from 3i using a 63x/1.4 NA Oil Plan-Apochromat DIC m27, and a 100x/1.46NA Oil Plan-Apochromat DIC m27 objective lens with 4X zoom via a SoRa Disc for super-resolution microscopy, respectively. All images were acquired from segments T3-A3 unless otherwise specified.

*For live preparations:* The following assay was used for live imaging of aCC/RP2 dendrites and/or astrocyte mitochondria in isolated CNS (Figures 1-2, Supplemental Movies 1-10). Larvae were dissected at 8 h ALH in a hemolymph-like solution (HL3.1); both lobes and the ventral nerve cord were kept intact. Isolated brains were mounted dorsal-side down on a 12mm #1 thickness poly-D-lysine coated round coverslips (Corning® BioCoat™, 354085 or German Glass Cover Slips, Fisher Scientific, 72196-12), a single 18mm x 18mm x 0.16mm cover glass (Fisher, 12-542B) was pressed against the coverslip until brain lobes were slightly compressed, a drop of HL3.1was used to facilitate sealing. The cover slip was then inverted onto a slide with vacuum grease as a bridge and pressed down until brain lobes were sandwiched between the slide and cover glass. Samples that were damaged during mounted were not imaged. Samples were imaged on a Zeiss LSM 800 laser scanning confocal fitted with a 63x/1.40 NA Oil Plan-Apochromat DIC m27 objective lens and GaAsP photomultiplier tubes, or on a spinning disc confocal from Intelligent Imaging Innovations (3i) using a 63x/1.4 NA Oil Plan-Apochromat DIC m27 objective lens, and imaged using a 488 nm laser and/or 561 nm laser when appropriate. For optogenetic experiments in Figures 1-2, both dissections and mounting were performed under low light conditions (<100 lx) to delay optogenetic activation. Continuous scans were obtained every 30 seconds for 15 minutes (z-stack of X µm, step of X µm to accommodate drift), with two hemisegments in the field of view. A z-stack of 12 µm (allowing for drift in Z) with .5 µm step size was performed. Finally, we imaged abdominal segments 5-6, which were left prone to drift over the 15-minute acquisition. For our mitochondria machinery screen (Supplemental Movies 5-8), we obtained 50 µm stacks (step size of 1 µm) once a minute for 15 minutes, with abdominal segments 5-8 in view. For GCaMP7s assays (Figure 7), we imaged a stereotyped .198 µm plan encompassing aCC/RP2 motor neuron cell bodies at 4 Hz for 5 minutes, with abdominal segments 1-6 in view.

### Behavioral analyses

We transferred larvae at either 8 h ALH or 72 h ALH from apple caps or food bottles (standard corn meal), respectively, to 1.2% agarose plates for half an hour prior to locomotion assays to allow the larvae to acclimate to the new crawling surface. Larvae were then transferred to a FIM behavior table (Risse et al., 2014) fitted with a fresh 1.2% agarose gel and allowed to further acclimate for two minutes prior to imaging. Two or more independent cohorts of larvae (N=30 larvae per cohort) were tested per genotype. Larval crawling was imaged at 4 Hz, 91 pixels/cm for one minute using a Basler acA2040-25gm camera in the Pylon5 Camera Software Suite (Basler). Data were then analyzed using FIMTrack software using standard settings (Risse et al., 2014). Data files were blinded to genotype until after analysis.

## QUANTIFICATION AND STATISTIC ANALYSIS

### Image processing and analyses

Quantitative analyses were performed either in Imaris (Bitplane 9.2.0 or later) or in FIJI (ImageJ 1.50d, https://imagej.net/Fiji).

### Synapse quantification: association of astrocyte mitochondria with pre-synapses

#### aCC/RP2-associated presynapses and mitochondrial volumes

For quantification of astrocyte mitochondria in close proximity to aCC/RP2-associated presynapses (Figures 1-2), data were acquired with a voxel size of .099 x .099 x .23 µm3 and de-convoluted in Imaris. Data for motor neuron mitochondria (Figures 3-4) were acquired with a voxel size of .060 x .060 x .270 µm^3^, and de-convoluted following the same protocol as with the astrocyte mitochondria. First, aCC/RP2 dendrites within a single hemisegment were reconstructed using the Imaris “Surface” module (no smoothing, thresholds varied with fluorescence intensity). A standard ROI spanned 100 x 100 pixels in XY, and 7 µm in Z with the top of the ROI beginning dorsally at the aCC/RP2 axons. The dendrite “Surface” was also used to determine dendrite volume. A distance transformation was then performed on the dendrite “Surface”. Second, Brp+ presynapses within the ROI were annotated using the Imaris “Spots” functions and then classified as verified post-synapses if they fell within 90 nm of the “Surface” (accounts for chromatic aberration and the size of the synaptic cleft) based on previously validated criteria (Ackerman et al., 2021; Sales et al., 2019), and were used to build a new channel. Third, astrocyte mitochondria within the same ROI were reconstructed using the Imaris “Surface” module (.1 µm smoothing, thresholds varied with fluorescence intensity). A distance transformation was then performed on the astrocyte mitochondria “Surface”. Finally, verified presynapse “Spots” were classified as mitochondria-associated if they fell within 120 nm of the astrocyte mitochondria “Surface” (accounts for chromatic aberration (Sales et al., 2019), the thickness of the astrocyte plasma membrane (Ingólfsson et al., 2017), and the thickness of both mitochondrial membranes (Perkins et al., 1997). For quantification of motor neuron mitochondria in close proximity to the same presynapses, the same method was following using a “Surface” reconstructed from motor neuron mitochondria fluorescence instead. Total mitochondrial “Surface” volume was provided by Imaris as an automatically generated value under the “Detailed” section of “Statistics”.

#### Astrocyte morphometry

For quantification of astrocyte membrane-associated presynapses (Figure 6), microtubule-based motor data were acquired with a voxel size of .2 x .2 x .39 µm^3^, and actin-based motor data were acquired with a voxel size of .060 x .060 x .270. Imaris reconstructions were applied to the entire larval brain. First, we reconstructed astrocyte membranes using the Imaris “Surface” module (no smoothing, thresholds varied with fluorescence intensity). A distance transformation was performed on the membrane “Surface.” Then, we constructed astrocyte mitochondria using the Imaris “Surface” module (no smoothing, thresholds varied with fluorescence intensity). A second distance transformation was performed on the mitochondria “Surface.” Finally, Brp+ presynapses were annotated using the Imaris “Spots” functions and then categorized by proximity (≤ 1 µm) to the astrocyte mitochondria or membrane “Surface.”

### Analysis of time-lapse imaging samples (optogenetics)

For analysis of mitochondrial dynamics over time (Figures 1-2), data were collected with a voxel size of .085 x .085 x .5 µm^3^. Any samples that drifted out of the frame of view during the acquisition period were not processed further. Data were then imported to Imaris and analyzed using the “Surface” module (.1 µm smoothing, thresholding varied by fluorescence intensity of sample). A standard ROI spanned 200 x 200 pixels in XY, and 12.5 µm in Z. First, we built a “Surface” to mask the motor dendrites. Then, we built a “Surface” to mask the astrocyte mitochondria. We then used the “Surface-Surface” overlap function (Imaris 9.7.0 or later) to calculate the volume of all mitochondria within the dendritic domain (≤ 120 nm from motor dendrites), and to determine the total volume of “Surface” overlap (*e.g.* volume of membrane overlap).

### Analysis of GCaMP imaging data

Image files were blinded to genotype. Raw time-lapse image stacks were imported into Fiji and converted to time-lapse maximum intensity projections. These converted files were loaded into a custom MATLAB script (Sales et al., 2019). The script first performs rigid registration to correct for movement artifacts during recording, and then allows for ROI selection. ROIs were drawn around aCC/RP2 dendrites in 6 individual hemisegments, 3 per side. ROI size was constant across all analyzed files (Figure 7). The script then quantifies and plots the frame-by-frame average dF/F within each ROI.

To identify peaks of calcium activity in an unbiased manner, our dF/F traces were next loaded into the MATLAB program PeakCaller (*Artimovich et al., 2017*). In PeakCaller, the following parameters were specified for the program to identify peaks in each trace: Required Rise = 20%, Required Fall = 30%, Max Lookback and Lookahead pts = 10 frames. Peaks < dF/F=0.1 were excluded from analysis. We used PeakCaller to identify the number of calcium peaks in each dF/F trace. We logged the amplitude of each peak (dF/F) within a trace, and averaged these amplitudes together to obtain a single average peak fluorescence value for each ROI.

After quantifying the frequency and amplitude of peaks for each ROI, sample files were unblinded, and statistical comparisons were performed between Control and *milton* RNAi groups using a two-tailed unpaired Student’s T-test.

### Analysis of synaptic boutons in larval fillet preparations

Data were acquired from dorsal muscles only (DO1-DO2, DA1-DA2) with a voxel size of.198 x .198 x .3 µm^3^. Samples were blinded and imported into Fiji, where we performed a max intensity projection of the channel corresponding to *vglut-lexA, lexAop-myr::GFP* (labels all motor neuron projections and synaptic boutons). Synaptic boutons were quantified manually using the Fiji “Multipoint” tool. After quantifying bouton numbers for an entire experimental set, sample files were unblinded and statistical comparisons were performed using a two-tailed unpaired Student’s T-test with unequal variance.

### Analysis of intercellular mitochondrial transfer

For quantification of astrocyte mitochondria within motor neuron membranes (Figure 5, Supplemental Figure 2), data were acquired with a voxel size of .027 x .027 x .015 µm^3^. Imaris reconstructions were applied to the entire larval brain. First, we reconstructed neuron membranes (Chrimson::TdTomato or GtACR2::EYFP+) T3-A3 using the Imaris “Surface” module (no smoothing, thresholds varied with fluorescence intensity). Then, we reconstructed astrocyte mitochondria within the same region of interest (ROI) once again using the Imaris “Surface” module (0.02 smoothing, thresholds varied with fluorescence intensity). Finally, we reconstructed astrocyte membrane (anti-GLUL) within that ROI using another Imaris “Surface” (no smoothing, thresholds varied with fluorescence intensity). Imaris provided the Surface-Surface overlap between these surfaces, which was used as a quantification of total volume of intercellularly transferred mitochondria. Mitochondrial “Surfaces” were examined by eye to determine which were entirely engulfed by neuronal membrane.

### Statistical analyses

Statistics were performed using a combination of Microsoft Excel, MATLAB (MathWorks), and Prism (GraphPad) software. We used a combination of one-way ANOVA, Student’s T-Tests (two-tailed, unequal variance), Kruskal-Wallis tests, and Mann-Whitney Tests where appropriate (noted throughout). Error bars, Standard Error of the Mean (SEM) unless otherwise noted. A 95% confidence interval was used to define the level of significance. Significance: *, p<.05; **, p<.01; ***, p<.001; ****, p<.0001, NS= not significant. All other pertinent information, including sample size, statistical test employed, and variance can be found in the figure legends or labeled within the figure.

## FIGURE PREPARATION

Images in figures were prepared in Imaris (Bitplane AG, version 9.2.0 or higher) or in Fiji (ImageJ 1.50d). Both software packages were used for rendering 3D, max, or orthogonal projections of the data, which were then assembled using Adobe Illustrator. Schematics were drawn in Microsoft Powerpoint.

## Supporting information

SupMovie1_RN2myrTdTomato_AlrmMitoGFP

SupMovie2_RN2Chrimson_AlrmMitoGFP

SupMovie3_RN2myrGFP_AlrmCherryMitoOMM

SupMovie4_RN2GtACR2_AlrmCherryMitoOMM

SupMovie5_ControlRNAi1

SupMovie6_MiltonRNAi

SupMovie7_Dhc64cRNAi

SupMovie8_DidumRNAi

SupplementalMovie9_ControlGCaMP

SupplementalMovie10_MiltonGCaMP

---

**Supplemental Figure 1.**
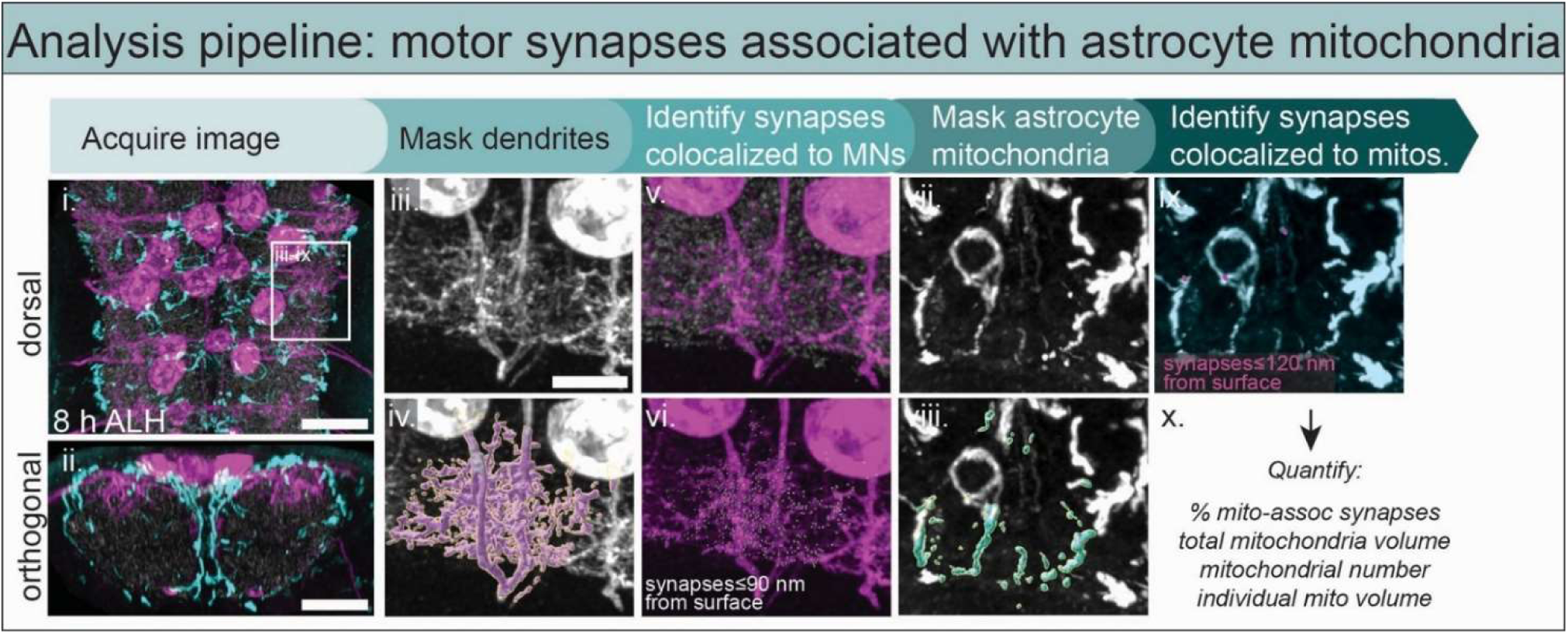
Analysis pipeline: motor synapses associated with astrocyte mitochondria. Pipeline for identifying motor synapses in close proximity to astrocyte mitochondria at 8 hours after larval hatching (h ALH). (**i-ii**) Acquire confocal z-stack through neuropil with fluorescently labeled astrocyte mitochondria (*R25H07-gal4, UAS-mito::GFP* or *alrm-lexA, lexAop-mcherry::mito.OMM*), aCC/RP2 motor neurons (MN; *RN2-lexA, lexAop-TdTomato* or *RN2-gal4, UAS-myr::GFP*), and presynapses (Brp+). Scale bar, 10 µm. (**iii-iv**) Mask the motor dendrites with an Imaris “Surface.” (**v-vi**) Identify presynapses colocalized with motor dendrites (≤90 nm) and generate Imaris “Spots”. (**vii-viii**) Mask astrocyte mitochondria with an Imaris “Surface.” (**ix-x**) Identify Spots within 120 nm of masked mitochondria. (**iii-ix**) Scale bar, 5 µm. For more details, see Methods.

**Supplemental Figure 2.**
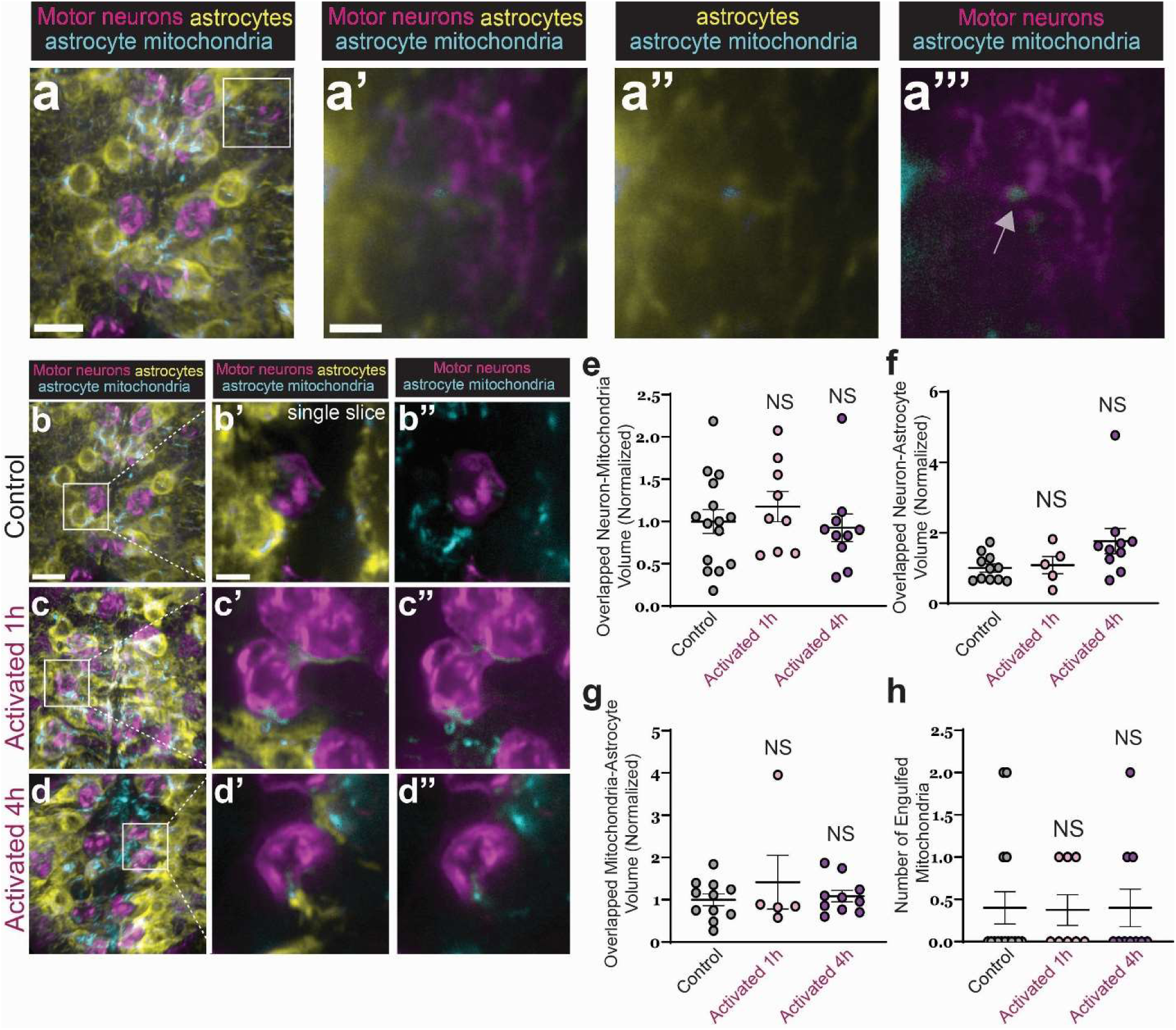
Intercellular transport of mitochondria from astrocytes to motor neurons is unaffected by ectopic neuronal activation. (**a-a’’’**) Representative images of astrocyte-neuron mitochondrial transfer in motor neuron dendrite. (a) Motor dendrites (magenta, *RN2-lexA, lexAop-Chrimson::tdTomato*), astrocyte membranes (yellow, GLUR+) with astrocyte mitochondria (cyan, *alrm-gal4, UAS-mito::GFP*) in fixed preparations at 8 h ALH. Prime panel shows a single 0.155 µm slice of neuron membrane, astrocyte membrane, and astrocyte mitochondria. Double prime panels show the same slice with only astrocyte and astrocyte mitochondria. Triple prime panels show the same slice with motor neuron dendrites and astrocyte mitochondria. Arrowhead denotes astrocyte mitochondria within motor neuron dendrite. (**a**) Scale bar, 5 µm. Prime panels: Scale bar 2, µm. (**b-d”**) Representative images of dark-reared control (**b-b’”**), 1-hour activated (**c-c’’**), and 4-hour activated (**d-d’’’**) motor dendrites (magenta, *RN2-lexA, lexAop-Chrimson::tdTomato*), astrocyte membranes (yellow, GLUL+) with astrocyte mitochondria (cyan, *alrm-gal4, UAS-mito::GFP*) in fixed preparations at 8 h ALH. Prime panels show a single 0.155 µm slice of astrocytes, neurons, and astrocyte mitochondria. Double prime panels show the same slice with motor neurons and astrocyte mitochondria. (**a,b,c**) Scale bar, 5 µm. Prime panels, Scale bar 2 µm. (e) Quantification of the neuron-astrocyte mitochondrial overlap Control: n=15 animals. 1-hour activated: n=9 animals (p>0.9999, Kruskal-Wallis test). 4-hour activated: n=10 (p>0.9999, Kruskal-Wallis test). (f) Quantification of astrocyte mitochondrial volume. Control: n=11 animals (p>0.9999, Kruskal-Wallis test). 1-h activated: n=5 animals (p>0.9999, Kruskal-Wallis test). 4-hour activated: n=10 animals (p=0.678, Kruskal-Wallis test). 119.8243 122.8806 (g) Quantification of the neuron-astroctye membrane overlap. Control: n=11 animals. 1-hour activated: n=5 animals (p>0.9999, Kruskal-Wallis test). 4-hour activated: n=10 animals (p>0.9999, Kruskal-Wallis test). (h) Count of astrocyte mitochondria found completely enveloped by neuron membrane. Control: n=15 animals. 1-hour activated: n=8 animals (p>0.9999, Kruskal-Wallis test). 4-hour activated: n=10 animals (p>0.9999, Kruskal-Wallis test). (**e-h**) NS, not significant. Error bars = SEM. N values reflect animals from 2 technical replicates.

**Supplemental Figure 3.**
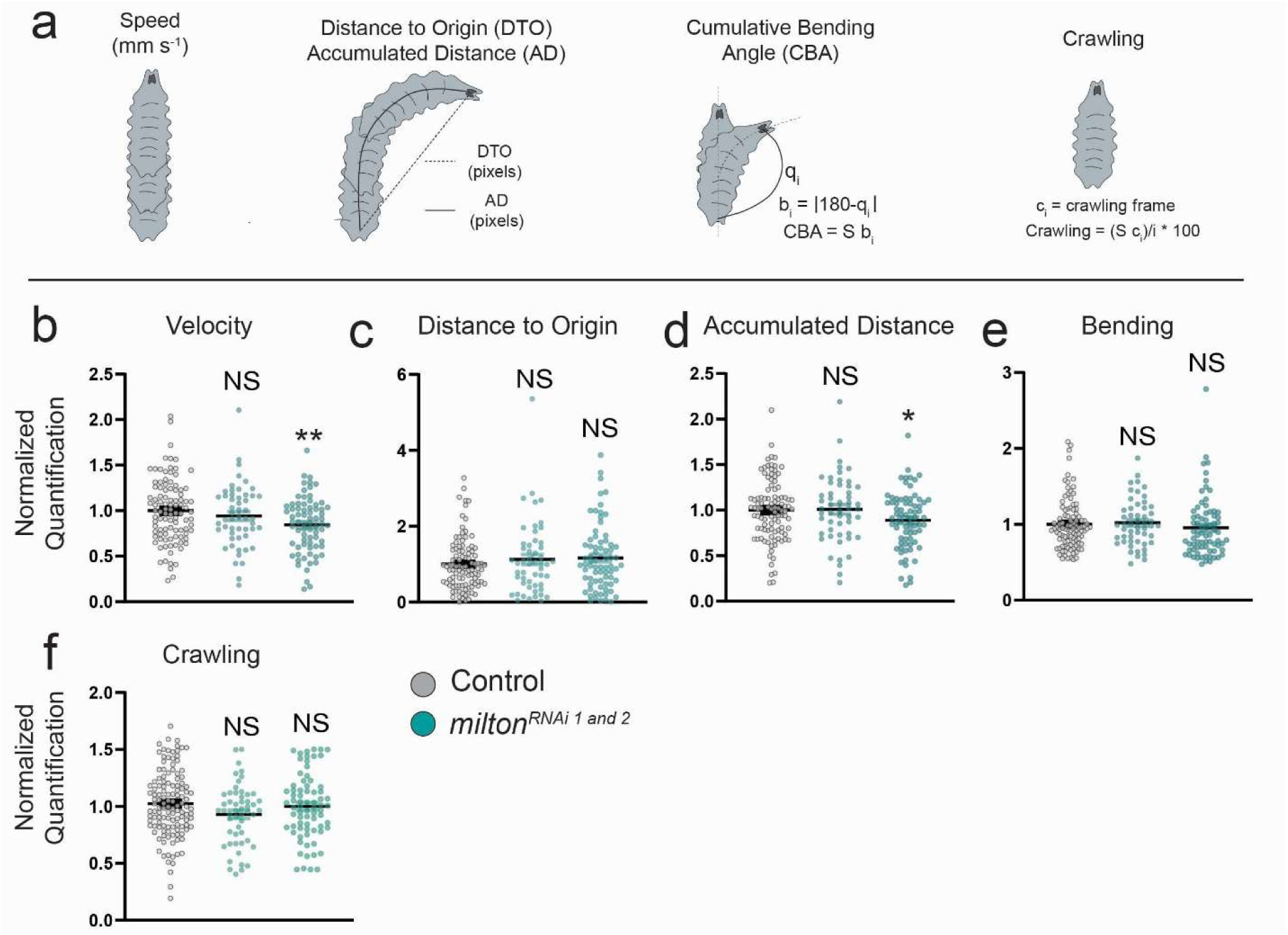
Loss of astrocyte peri-synaptic mitochondria causes subtle developmental behavioral defects. **(a)** Schematic of behavioral paradigms tested at 8 h ALH. **(b-f)** FIMTrack traces from individual, free-crawling larvae recorded for 2 minutes at 4 Hz*. luciferase^RNAi^* (luc) Control: n=100 animals. *milton^RNAi1^*: n=52 animals. *milton^RNAi2^*: n=73 animals. N values reflect animals from 3 technical replicates. **(g-k)** Quantification of behavioral metrics relative to control. Statistical significance determined by Mann-Whitney U Test for (**f**) velocity (*milton^RNAi1^*, p<.42; *milton^RNAi2^*, p<.01), (**g**) distance to origin (*milton^RNAi1^*, p<.62; *milton*^RNAi2^, p<.32), (**h**) accumulated distance (*milton^RNAi1^*, p<.87; *milton^RNAi2^*, p<.05), **(i)** cumulative bending angle (*milton^RNAi1^*, p<.49; *milton^RNAi2^*, p<.16), and (**j**) crawling (*milton^RNAi1^*, p<.6; *milton^RNAi2^*, p<.08). Error bars = SEM. NS, not significant.

**Supplemental Figure 4.**
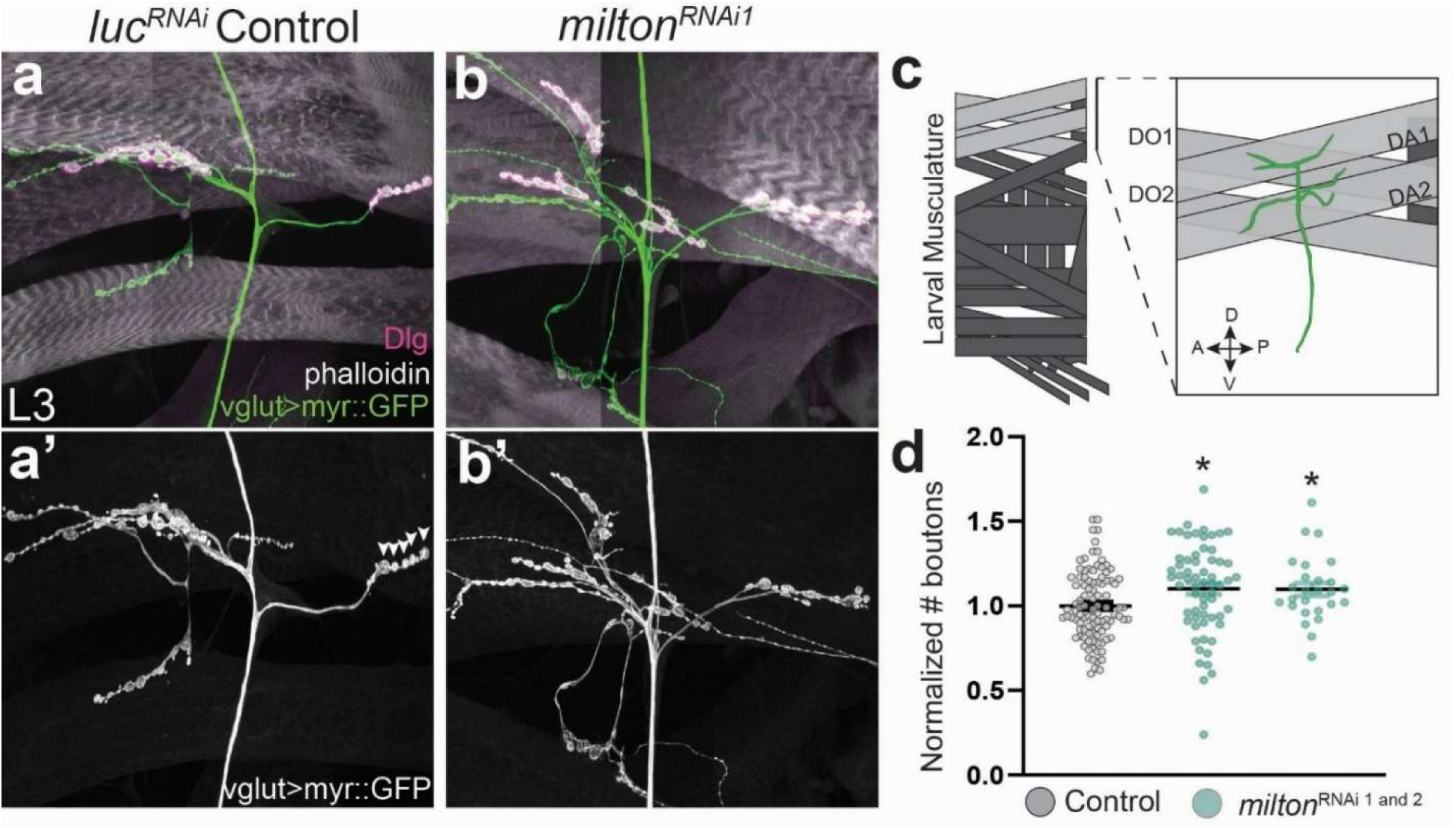
Astrocytic *milton* in the CNS modulates NMJ synapse number. **(a-b’)** Representative images of L3 neuromuscular junctions (NMJs, Green: *vglut-lexA, lexAop-myr::GFP*; Magenta: Dlg+; White: Phalloidin) from *luciferase^RNAi^* (luc) control (**a**) or following astrocyte-specific knockdown of *milton* (*R25H07-gal4, UAS-milton^RNAi^*). Prime panels show GFP channel used for quantification of synaptic boutons (arrowheads). Control: n=98 NMJs from n=16 animals. *milton^RNAi1^*: n=61 NMJs from n=12 animals. *milton^RNAi2^*: n=25 NMJs from n=4 animals. N values reflect animals from 2 technical replicates. (**c**) Schematic for reader orientation. (**d**) Quantification of bouton number (type 1b only). Statistical significance determined by Student’s T-Test relative to control: *milton^RNAi1^* (p<.02), *milton^RNAi2^* (p<.03).

## Supplemental Movie Legends

**Supplemental Movie 1: Time-lapse imaging of astrocyte mitochondria in myr::TdTomato control for Chrimson-activation.** Representative time-lapse movie showing astrocyte mitochondrial dynamics (*R25H07-gal4, UAS-mito::GFP*, cyan) around aCC/RP2 dendrites (*RN2-lexA, lexAop-myr::TdTomato*, magenta) in a fictive brain preparation at 8 h ALH. Animals were supplied with ATR and reared in the dark to mimic conditions for Chrimson-activation. A single z-stack was acquired every 30 seconds (each stack taking 25” to acquire) and brains were imaged for a total of 15 minutes. Astrocyte mitochondria were highly motile, though largely stable in volume over the 15-minute period. Left hemisegment shows Imaris “Surface” reconstruction of astrocyte mitochondria, right shows raw data in neighboring hemisegment.

**Supplemental Movie 2: Time-lapse imaging of astrocyte mitochondria during motor neuron Chrimson-activation.** Representative time-lapse movie showing astrocyte mitochondrial dynamics (*R25H07-gal4, UAS-mito::GFP*, cyan) around aCC/RP2 dendrites (*RN2-lexA, lexAop-Chrimson::TdTomato*, magenta) in a fictive brain preparation at 8 h ALH. Animals were supplied with ATR and dark-reared. A single z-stack was acquired every 30 seconds (each stack taking 25” to acquire) and brains were imaged for a total of 15 minutes. Astrocyte mitochondria were rapidly trafficked towards motor dendrites in the first several minutes, followed by relative stability. Left hemisegment shows Imaris “Surface” reconstruction of astrocyte mitochondria, right shows raw data in neighboring hemisegment.

**Supplemental Movie 3. Time-lapse imaging of astrocyte mitochondria in myr::GFP control for GtACR2 silencing.** Representative time-lapse movie showing astrocyte mitochondrial dynamics (*alrm-lexA, lexAop-mcherry::mito.OMM*, cyan) around aCC/RP2 dendrites (*RN2-gal4, UAS-GtACR2::EYFP*, magenta) in a fictive brain preparation at 8 h ALH. Animals were supplied with ATR and reared in the dark to mimic conditions for GtACR2-silencing. A single z-stack was acquired every 30 seconds (each stack taking 25” to acquire) and brains were imaged for a total of 15 minutes. Astrocyte mitochondria were highly motile, though largely stable in volume over the 15-minute period. Left hemisegment shows Imaris “Surface” reconstruction of astrocyte mitochondria, right shows raw data in neighboring hemisegment.

**Supplemental Movie 4. Time-lapse imaging of astrocyte mitochondria during motor neuron GtACR2-silencing.** Representative time-lapse movie showing astrocyte mitochondrial dynamics (*alrm-lexA, lexAop-mcherry::mito.OMM*, cyan) around aCC/RP2 dendrites (*RN2-gal4, UAS-GtACR2::EYFP*, magenta) in a fictive brain preparation at 8 h ALH. Animals were supplied with ATR and reared in the dark. A single z-stack was acquired every 30 seconds (each stack taking 25” to acquire) and brains were imaged for a total of 15 minutes. Astrocyte mitochondria were highly motile, though largely stable in volume over the 15-minute period. Left hemisegment shows Imaris “Surface” reconstruction of astrocyte mitochondria, right shows raw data in neighboring hemisegment.

**Supplemental Movie 5. Time lapse imaging showing control astrocyte mitochondrial dynamics.** Representative time-lapse movie showing astrocyte mitochondria (*R25H07-gal4, UAS-mito::GFP*) dynamics in a control brain (*UAS-luciferase^RNAi^*) at 8 h ALH. A single z-stack was acquired one a minute (each stack taking 15” to acquire) and brains were imaged for a total of 10 minutes. Mitochondria were distributed throughout the neuropil.

**Supplemental Movie 6. Time lapse imaging showing astrocyte mitochondrial dynamics following astrocyte knockdown of *milton*.** Representative time-lapse movie showing astrocyte mitochondria (*R25H07-gal4, UAS-mito::GFP*) dynamics in a milton knockdown brain (*UAS-milton^RNAi^*) at 8 h ALH. A single z-stack was acquired one a minute (each stack taking 15” to acquire) and brains were imaged for a total of 10 minutes. Mitochondria were absent from the neuropil.

**Supplemental Movie 7. Time lapse imaging showing astrocyte mitochondrial dynamics following astrocyte knockdown of *Dhc64C*.** Representative time-lapse movie showing astrocyte mitochondria (*R25H07-gal4, UAS-mito::GFP*) dynamics in a *Dhc64C* knockdown brain (*UAS-Dhc64C^RNAi^*) at 8 h ALH. A single z-stack was acquired one a minute (each stack taking 15” to acquire) and brains were imaged for a total of 10 minutes. Mitochondria distributed normally in the neuropil.

**Supplemental Movie 8. Time lapse imaging showing astrocyte mitochondrial dynamics following astrocyte knockdown of *jar*.** Representative time-lapse movie showing astrocyte mitochondria (*R25H07-gal4, UAS-mito::GFP*) dynamics in a *jar* knockdown brain (*UAS-jar^RNAi^*) at 8 h ALH. A single z-stack was acquired one a minute (each stack taking 15” to acquire) and brains were imaged for a total of 10 minutes. Mitochondria distributed less abundantly in the neuropil.

**Supplemental Movie 9. Time-lapse imaging of control aCC/RP2 activity.** Representative time-lapse movie (40 fps) showing control aCC/RP2 activity in a fictive brain preparation at 8 h ALH (*RN2-lexA, lexAop-GCaMP7s*; *R25H07-gal4, UAS-luciferase^RNAi^*). A single slice was acquired every 250 ms and brains were imaged for a total of 5 minutes.

**Supplemental Movie 10. Time-lapse imaging of aCC/RP2 activity following astrocyte knockdown of** *milton*. Representative time-lapse movie (40 fps) showing aCC/RP2 activity in a fictive brain preparation at 8 h ALH (*RN2-lexA, lexAop-GCaMP7s*) following astrocyte knockdown of *milton* (*R25H07-gal4, UAS-milton*^RNAi^). A single slice was acquired every 250 ms and brains were imaged for a total of 5 minutes. Activity was reduced relative to control.

## Notes

### Competing Interest Statement

The authors have declared no competing interest.

### Summary of Updates

This manuscript was revised to update all associated figures.

### Funding

- NIH: 1R01NS144659

## References

1. Ackerman, S. D., Perez-Catalan, N. A., Freeman, M. R., & Doe, C. Q. (2021b). Astrocytes close a motor circuit critical period. Nature, 592(7854), 414–420. 10.1038/s41586-021-03441-2

2. Allen, N. J., & Eroglu, C. (2017). Cell Biology of Astrocyte-Synapse Interactions. Neuron, 96(3), 697– 708. 10.1016/j.neuron.2017.09.056

3. Annesley, S. J., & Fisher, P. R. (2019). Mitochondria in Health and Disease. Cells, 8(7), 680. 10.3390/cells8070680

4. Artimovich: PeakCaller: An automated graphical interface… – Google Scholar. (n.d.). Retrieved July 15, 2025, from https://scholar.google.com/scholar_lookup?author=E.+Artimovich&author=R.+K.+Jackson&author=M.+B.+C.+Kilander&author=Y.-C.+Lin&author=M.+W+Nestor&title=PeakCaller%3A+an+automated+graphical+interface+for+the+quantification+of+intracellular+calcium+obtained+by+high-content+screening&publication_year=2017&journal=BMC+Neurosci&volume=18

5. Aten, S., Kiyoshi, C. M., Arzola, E. P., Patterson, J. A., Taylor, A. T., Du, Y., Guiher, A. M., Philip, M., Camacho, E. G., Mediratta, D., Collins, K., Boni, K., Garcia, S. A., Kumar, R., Drake, A. N., Hegazi, A., Trank, L., Benson, E., Kidd, G., … Zhou, M. (2022). Ultrastructural view of astrocyte arborization, astrocyte-astrocyte and astrocyte-synapse contacts, intracellular vesicle-like structures, and mitochondrial network. Progress in Neurobiology, 213, 102264. 10.1016/j.pneurobio.2022.102264

6. Attwell, D., & Laughlin, S. B. (2001). An energy budget for signaling in the grey matter of the brain. Journal of Cerebral Blood Flow and Metabolism: Official Journal of the International Society of Cerebral Blood Flow and Metabolism, 21(10), 1133–1145. 10.1097/00004647-200110000-00001

7. Balasubramanian, V. (2021). Brain power. Proceedings of the National Academy of Sciences, 118(32). 10.1073/pnas.2107022118

8. Bastian, C., Zerimech, S., Nguyen, H., Doherty, C., Franke, C., Faris, A., Quinn, J., & Baltan, S. (2022). Aging astrocytes metabolically support aging axon function by proficiently regulating astrocyte-neuron lactate shuttle. Experimental Neurology, 357, 114173. 10.1016/j.expneurol.2022.114173

9. Bélanger, M., Allaman, I., & Magistretti, P. J. (2011). Brain energy metabolism: Focus on astrocyte-neuron metabolic cooperation. Cell Metabolism, 14(6), 724–738. 10.1016/j.cmet.2011.08.016

10. Benjamin Kacerovsky, J., & Murai, K. K. (2016). Stargazing: Monitoring subcellular dynamics of brain astrocytes. Neuroscience, 323, 84–95. 10.1016/j.neuroscience.2015.07.007

11. Berridge, M. V., & Neuzil, J. (2017). The mobility of mitochondria: Intercellular trafficking in health and disease. Clinical and Experimental Pharmacology & Physiology, 44 *Suppl 1*, 15–20. 10.1111/1440-1681.12764

12. Birsa, N., Norkett, R., Higgs, N., Lopez-Domenech, G., & Kittler, J. T. (2013). Mitochondrial trafficking in neurons and the role of the Miro family of GTPase proteins. Biochemical Society Transactions, 41(6), 1525–1531. 10.1042/BST20130234

13. Bonvento, G., & Bolaños, J. P. (2021). Astrocyte-neuron metabolic cooperation shapes brain activity. Cell Metabolism, 33(8), 1546–1564. 10.1016/j.cmet.2021.07.006

14. Brickley, K., & Stephenson, F. A. (2011). Trafficking kinesin protein (TRAK)-mediated transport of mitochondria in axons of hippocampal neurons. The Journal of Biological Chemistry, 286(20), 18079–18092. 10.1074/jbc.M111.236018

15. Chandel, N. S. (2021). Glycolysis. Cold Spring Harbor Perspectives in Biology, 13(5), a040535. 10.1101/cshperspect.a040535

16. Course, M. M., & Wang, X. (2016). Transporting mitochondria in neurons. F1000 Research, 5, F1000 Faculty Rev-1735. 10.12688/f1000research.7864.1

17. Crisp: The development of motor coordination in Drosophil… – Google Scholar. (n.d.). Retrieved July 15, 2025, from https://scholar.google.com/scholar_lookup?author=S.+Crisp&author=J.+F.+Evers&author=A.+Fiala&author=M+Bate&title=The+development+of+motor+coordination+in+Drosophila+embryos&publication_year=2008&journal=Dev.+Camb.+Engl&volume=135&pages=3707-3717

18. Doherty: Ensheathing glia function as phagocytes… – Google Scholar. (n.d.). Retrieved July 15, 2025, from https://scholar.google.com/scholar_lookup?author=J.+Doherty&author=M.+A.+Logan&author=O.+E.+Ta%C5%9Fdemir&author=M.+R+Freeman&title=Ensheathing+glia+function+as+phagocytes+in+the+adult+Drosophila+brain&publication_year=2009&journal=J.+Neurosci.+Off.+J.+Soc.+Neurosci&volume=29&pages=4768-4781

19. Freeman, M. R. (2010). Specification and morphogenesis of astrocytes. Science (New York, N.Y.), 330(6005), 774–778. 10.1126/science.1190928

20. Glater, E. E., Megeath, L. J., Stowers, R. S., & Schwarz, T. L. (2006). Axonal transport of mitochondria requires milton to recruit kinesin heavy chain and is light chain independent. The Journal of Cell Biology, 173(4), 545–557. 10.1083/jcb.200601067

21. Govorunova, E. G., Sineshchekov, O. A., Janz, R., Liu, X., & Spudich, J. L. (2015). Natural light-gated anion channels: A family of microbial rhodopsins for advanced optogenetics. Science, 349(6248), 647–650. 10.1126/science.aaa7484

22. Hall, K. D., Heymsfield, S. B., Kemnitz, J. W., Klein, S., Schoeller, D. A., & Speakman, J. R. (2012). Energy balance and its components: Implications for body weight regulation. The American Journal of Clinical Nutrition, 95(4), 989–994. 10.3945/ajcn.112.036350

23. Harman, D. (1972). The biologic clock: The mitochondria? Journal of the American Geriatrics Society, 20(4), 145–147. 10.1111/j.1532-5415.1972.tb00787.x

24. Hayakawa, K., Esposito, E., Wang, X., Terasaki, Y., Liu, Y., Xing, C., Ji, X., & Lo, E. H. (2016). Transfer of mitochondria from astrocytes to neurons after stroke. Nature, 535(7613), 551–555. 10.1038/nature18928

25. Heo, J.-M., Ordureau, A., Paulo, J. A., Rinehart, J., & Harper, J. W. (2015). The PINK1-PARKIN Mitochondrial Ubiquitylation Pathway Drives a Program of OPTN/NDP52 Recruitment and TBK1 Activation to Promote Mitophagy. Molecular Cell, 60(1), 7–20. 10.1016/j.molcel.2015.08.016

26. Hutto, R. A., Rutter, K. M., Giarmarco, M. M., Parker, E. D., Chambers, Z. S., & Brockerhoff, S. E. (2023). Cone photoreceptors transfer damaged mitochondria to Müller glia. Cell Reports, 42(2), 112115. 10.1016/j.celrep.2023.112115

27. Ingólfsson: Computational lipidomics of the neuronal… – Google Scholar. (n.d.). Retrieved July 15, 2025, from https://scholar.google.com/scholar_lookup?author=H.+I.+Ing%C3%B3lfsson&title=Computational+Lipidomics+of+the+Neuronal+Plasma+Membrane&publication_year=2017&journal=Biophys&volume=113&pages=2271-2280

28. Jackson, J. G., O’Donnell, J. C., Takano, H., Coulter, D. A., & Robinson, M. B. (2014). Neuronal activity and glutamate uptake decrease mitochondrial mobility in astrocytes and position mitochondria near glutamate transporters. The Journal of Neuroscience: The Official Journal of the Society for Neuroscience, 34(5), 1613–1624. 10.1523/JNEUROSCI.3510-13.2014

29. Jackson, J. G., & Robinson, M. B. (2015). Reciprocal Regulation of Mitochondrial Dynamics and Calcium Signaling in Astrocyte Processes. The Journal of Neuroscience: The Official Journal of the Society for Neuroscience, 35(45), 15199–15213. 10.1523/JNEUROSCI.2049-15.2015

30. Joshi, A. U., Minhas, P. S., Liddelow, S. A., Haileselassie, B., Andreasson, K. I., Dorn II, G. W., & Mochly-Rosen, D. (2019). Fragmented mitochondria released from microglia trigger A1 astrocytic response and propagate inflammatory neurodegeneration. Nature Neuroscience, 22(10), 1635–1648. 10.1038/s41593-019-0486-0

31. Kann, M. R., Ackerman, M. K., & Ackerman, S. D. (2024). OptoChamber: A Low-cost, Easy-to-Make, Customizable, and Multi-Chambered Electronic Device for Applying Optogenetic Stimulation to Larval Drosophila melanogaster. microPublication Biology, 2024. 10.17912/micropub.biology.001082

32. Kim, N.-S., & Chung, W.-S. (2023). Astrocytes regulate neuronal network activity by mediating synapse remodeling. Neuroscience Research, 187, 3–13. 10.1016/j.neures.2022.09.007

33. Klapoetke, N. C., Murata, Y., Kim, S. S., Pulver, S. R., Birdsey-Benson, A., Cho, Y. K., Morimoto, T. K., Chuong, A. S., Carpenter, E. J., Tian, Z., Wang, J., Xie, Y., Yan, Z., Zhang, Y., Chow, B. Y., Surek, B., Melkonian, M., Jayaraman, V., Constantine-Paton, M., … Boyden, E. S. (2014). Independent optical excitation of distinct neural populations. Nature Methods, 11(3), 338–346. 10.1038/nmeth.2836

34. Lago-Baldaia, I., Fernandes, V. M., & Ackerman, S. D. (2020). More Than Mortar: Glia as Architects of Nervous System Development and Disease. Frontiers in Cell and Developmental Biology, 8, 611269. 10.3389/fcell.2020.611269

35. Landgraf, M., Jeffrey, V., Fujioka, M., Jaynes, J. B., & Bate, M. (2003). Embryonic Origins of a Motor System: Motor Dendrites Form a Myotopic Map in Drosophila. PLOS Biology, 1(2), e41. 10.1371/journal.pbio.0000041

36. Lardy, H. A., & Ferguson, S. M. (1969). Oxidative phosphorylation in mitochondria. Annual Review of Biochemistry, 38, 991–1034. 10.1146/annurev.bi.38.070169.005015

37. Magistretti, P. J., Pellerin, L., Rothman, D. L., & Shulman, R. G. (1999). Energy on Demand. Science, 283(5401), 496–497. 10.1126/science.283.5401.496

38. Malik, D. M., Rhoades, S. D., Kain, P., Sengupta, A., Sehgal, A., & Weljie, A. M. (2024). Altered Metabolism during the Dark Period in Drosophila Short Sleep Mutants. Journal of Proteome Research, 23(9), 3823–3836. 10.1021/acs.jproteome.4c00106

39. Mallik, B., & Frank, C. A. (2022). Roles for Mitochondrial Complex I Subunits in Regulating Synaptic Transmission and Growth. Frontiers in Neuroscience, 16. 10.3389/fnins.2022.846425

40. Mauss: Optogenetic neuronal silencing in Drosophila… – Google Scholar. (n.d.). Retrieved July 15, 2025, from https://scholar.google.com/scholar_lookup?author=A.+S.+Mauss&author=C.+Busch&author=A+Borst&title=Optogenetic+Neuronal+Silencing+in+Drosophila+during+Visual+Processing&publication_year=2017&journal=Sci.+Rep&volume=7

41. Melkov, A., & Abdu, U. (2018). Regulation of long-distance transport of mitochondria along microtubules. Cellular and Molecular Life Sciences: CMLS, 75(2), 163–176. 10.1007/s00018-017-2590-1

42. Melkov, A., Baskar, R., Alcalay, Y., & Abdu, U. (2016). A new mode of mitochondrial transport and polarized sorting regulated by Dynein, Milton and Miro. *Development (Cambridge*, England), 143(22), 4203–4213. 10.1242/dev.138289

43. Nath, S., & Villadsen, J. (2015). Oxidative phosphorylation revisited. Biotechnology and Bioengineering, 112(3), 429–437. 10.1002/bit.25492

44. Oberheim, N. A., Takano, T., Han, X., He, W., Lin, J. H. C., Wang, F., Xu, Q., Wyatt, J. D., Pilcher, W., Ojemann, J. G., Ransom, B. R., Goldman, S. A., & Nedergaard, M. (2009). Uniquely hominid features of adult human astrocytes. The Journal of Neuroscience: The Official Journal of the Society for Neuroscience, 29(10), 3276–3287. 10.1523/JNEUROSCI.4707-08.2009

45. Peco: Drosophila astrocytes cover specific territories… – Google Scholar. (n.d.). Retrieved July 15, 2025, from https://scholar.google.com/scholar_lookup?author=E.+Peco&title=Drosophila+astrocytes+cover+specific+territories+of+the+CNS+neuropil+and+are+instructed+to+differentiate+by+Prospero%2C+a+key+effector+of+Notch&publication_year=2016&journal=Dev.+Camb.+Engl&volume=143&pages=1170-1181

46. Pellerin, L., & Magistretti, P. J. (1994). Glutamate uptake into astrocytes stimulates aerobic glycolysis: A mechanism coupling neuronal activity to glucose utilization. Proceedings of the National Academy of Sciences of the United States of America, 91(22), 10625–10629. 10.1073/pnas.91.22.10625

47. Perkins: Electron tomography of neuronal mitochondria:… – Google Scholar. (n.d.). Retrieved July 15, 2025, from https://scholar.google.com/scholar_lookup?author=G.+Perkins&title=Electron+tomography+of+neuronal+mitochondria%3A+three-dimensional+structure+and+organization+of+cristae+and+membrane+contacts&publication_year=1997&journal=J.+Struct.+Biol&volume=119&pages=260-272

48. Pilling, A. D., Horiuchi, D., Lively, C. M., & Saxton, W. M. (2006). Kinesin-1 and Dynein are the primary motors for fast transport of mitochondria in Drosophila motor axons. Molecular Biology of the Cell, 17(4), 2057–2068. 10.1091/mbc.e05-06-0526

49. Reczek, C. R., & Chandel, N. S. (2015). ROS-dependent signal transduction. Current Opinion in Cell Biology, 33, 8–13. 10.1016/j.ceb.2014.09.010

50. Risse, B., Otto, N., Berh, D., Jiang, X., & Klämbt, C. (2014). FIM imaging and FIMtrack: two new tools allowing high-throughput and cost effective locomotion analysis. Journal of visualized experiments: JoVE, (94), 52207. 10.3791/52207

51. Sales, E. C., Heckman, E. L., Warren, T. L., & Doe, C. Q. (2019). Regulation of subcellular dendritic synapse specificity by axon guidance cues. eLife, 8, e43478. 10.7554/eLife.43478

52. Schwarz, T. L. (2013). Mitochondrial trafficking in neurons. Cold Spring Harbor Perspectives in Biology, 5(6), a011304. 10.1101/cshperspect.a011304

53. Senior, A. E. (1988). ATP synthesis by oxidative phosphorylation. Physiological Reviews, 68(1), 177–231. 10.1152/physrev.1988.68.1.177

54. Sheng, Z.-H., & Cai, Q. (2012). Mitochondrial transport in neurons: Impact on synaptic homeostasis and neurodegeneration. Nature Reviews. Neuroscience, 13(2), 77–93. 10.1038/nrn3156

55. Smith, H. L., Bourne, J. N., Cao, G., Chirillo, M. A., Ostroff, L. E., Watson, D. J., & Harris, K. M. (2016). Mitochondrial support of persistent presynaptic vesicle mobilization with age-dependent synaptic growth after LTP. eLife, 5, e15275. 10.7554/eLife.15275

56. Stork, T., Sheehan, A., Tasdemir-Yilmaz, O. E., & Freeman, M. R. (2014). Neuron-glia interactions through the Heartless FGF receptor signaling pathway mediate morphogenesis of Drosophila astrocytes. Neuron, 83(2), 388–403. 10.1016/j.neuron.2014.06.026

57. Suzuki, A., Stern, S. A., Bozdagi, O., Huntley, G. W., Walker, R. H., Magistretti, P. J., & Alberini, C. M. (2011). Astrocyte-Neuron Lactate Transport Is Required for Long-Term Memory Formation. Cell, 144(5), 810–823. 10.1016/j.cell.2011.02.018

58. Theisen, E. K., Rivas-Serna, I. M., Lee, R. J., Jay, T. R., Kunduri, G., Nguyen, T. T., Freeman, M. R., Mazurak, V., Clandinin, M. T., & Vaughen, J. P. Glia phagocytose neuronal sphingolipids to infiltrate developing synapses.

59. Theisen, E. K., Rivas-Serna, I. M., Lee, R. J., Jay, T. R., Kunduri, G., Nguyen, T. T., Mazurak, V., Clandinin, M. T., Clandinin, T. R., & Vaughen, J. P. (2025). Glia phagocytose neuronal sphingolipids to infiltrate developing synapses. bioRxiv: The Preprint Server for Biology, 2025.04.14.648777. 10.1101/2025.04.14.648777

60. Tsacopoulos, M., & Magistretti, P. J. (1996). Metabolic coupling between glia and neurons. The Journal of Neuroscience: The Official Journal of the Society for Neuroscience, 16(3), 877–885. 10.1523/JNEUROSCI.16-03-00877.1996

61. Tseng, N., Lambie, S. C., Huynh, C. Q., Sanford, B., Patel, M., Herson, P. S., & Ormond, D. R. (2021). Mitochondrial transfer from mesenchymal stem cells improves neuronal metabolism after oxidant injury in vitro: The role of Miro1. Journal of Cerebral Blood Flow and Metabolism: Official Journal of the International Society of Cerebral Blood Flow and Metabolism, 41(4), 761– 770. 10.1177/0271678X20928147

62. Wang, Z., Ying, Z., Bosy-Westphal, A., Zhang, J., Schautz, B., Later, W., Heymsfield, S. B., & Müller, M. J. (2010). Specific metabolic rates of major organs and tissues across adulthood: Evaluation by mechanistic model of resting energy expenditure. The American Journal of Clinical Nutrition, 92(6), 1369–1377. 10.3945/ajcn.2010.29885

63. Wu, Y., Ding, C., Sharif, B., Weinreb, A., Swaim, G., Hao, H., Yogev, S., Watanabe, S., & Hammarlund, M. (2024). Polarized localization of kinesin-1 and RIC-7 drives axonal mitochondria anterograde transport. The Journal of Cell Biology, 223(5), e202305105. 10.1083/jcb.202305105

64. Yates, D. (2024). A neuronal subcompartment view of ATP production. Nature Reviews Neuroscience, 25(3), 142–142. 10.1038/s41583-023-00792-9

65. Yellen, G. (2018). Fueling thought: Management of glycolysis and oxidative phosphorylation in neuronal metabolism. The Journal of Cell Biology, 217(7), 2235–2246. 10.1083/jcb.201803152

66. Zhou, J., Zhang, L., Peng, J., Zhang, X., Zhang, F., Wu, Y., Huang, A., Du, F., Liao, Y., He, Y., Xie, Y., Gu, L., Kuang, C., Ou, W., Xie, M., Tu, T., Pang, J., Zhang, D., Guo, K., Feng, Y., … Jiang, Y. (2024). Astrocytic LRP1 enables mitochondria transfer to neurons and mitigates brain ischemic stroke by suppressing ARF1 lactylation. Cell metabolism, 36(9), 2054–2068.e14. 10.1016/j.cmet.2024.05.016

67. Zhu, W., Miao, Q., Sun, D., Yang, G., Wu, C., Huang, J., & Zheng, C. (2012). The Mitochondrial Phosphate Transporters Modulate Plant Responses to Salt Stress via Affecting ATP and Gibberellin Metabolism in Arabidopsis thaliana. PLoS ONE, 7(8), e43530. 10.1371/journal.pone.0043530

